# Convergent Decay of Skin-specific Gene Modules in Pangolins

**DOI:** 10.1101/2022.12.08.519613

**Authors:** Bernardo Pinto, Raul Valente, Filipe Caramelo, Raquel Ruivo, L. Filipe C. Castro

**Affiliations:** CIMAR/CIIMAR - Interdisciplinary Centre of Marine and Environmental Research, University of Porto, Avenida General Norton de Matos, S/N, 4450-208 Matosinhos, Portugal; FCUP - Department of Biology, Faculty of Sciences, University of Porto (U. Porto), Rua do Campo Alegre, Porto, Portugal

**Keywords:** gene loss, skin, sebum, sweat glands, immunity

## Abstract

The Mammalia skin exhibits a rich spectrum of evolutionary adaptations. The pilosebaceous unit, composed of the hair shaft, follicle, and the sebaceous gland, is the most striking synapomorphy. The evolutionary diversification of mammals across different ecological niches was paralleled by the appearance of an ample variety of skin modifications. Pangolins, order Pholidota, exhibit keratin-derived scales, one of the most iconic skin appendages. This formidable armor is intended to serve as a deterrent against predators. Surprisingly, while pangolins have hair on their abdomens, the occurrence of sebaceous and sweat glands is contentious. Here, we explore various molecular modules of skin physiology in four pangolin genomes, including that of sebum production. We show that genes driving wax monoester formation - *Awat1/2*, show patterns of inactivation in the stem pangolin branch; while the triacylglycerol synthesis gene *Dgat2l6* seems independently eroded in the African and Asian clades. In contrast, *Elovl3* implicated in the formation of specific neutral lipids required for skin barrier function, is intact and expressed in the skin. An extended comparative analysis shows that genes involved in skin pathogen defense and structural integrity of keratinocyte layers also show inactivating mutations: associated with both ancestral and independent pseudogenization events. Finally, we deduce that the suggested absence of sweat glands is not paralleled by the inactivation of the ATP-binding cassette transporter *Abcc11*, as previously described in Cetacea. Our findings reveal the sophisticated, convergent and complex history of gene retention and loss as key mechanisms in the evolution of the highly modified mammalian skin phenotypes.

## Introduction

The evolution of mammals entailed some tantalizing lifestyle variations. Ecological transitions such as subterranean burrowing, powered flight or obligate aquatic regimes, elaborated from prominent eco-physiological adaptations, notably in the skin (Themudo et al. 2020; Wu et al. 2022). Some of these skin-phenotypic shifts were quite radical, as illustrated by the complete absence of glands and pelage in Cetacea skin (Figure 1). The molecular foundations underscoring the skin phenotype of Cetacea is contingent on gene repertoire variations (Nery et al. 2014; Springer and Gatesy 2018; Lopes-Marques et al. 2019; Springer et al. 2020; Themudo et al. 2020; Kowalczyk et al. 2021; Holthaus et al. 2021; Fuchs et al. 2022), which translate into a thick and smooth skin, to counterbalance the mechanical and thermal stress associated with an obligatory aquatic lifestyle (Spearman 1972; Reeb et al. 2007). In other mammalian lineages, the morphological co-occurrence of hair and associated glands (i.e. the pilosebaceous unit), has been more challenging to ascertain. In manatees (e.g., *Trichechus latirostris*), for instance, complete sebaceous gland regression is still disputed, while in the semi-aquatic hippopotamus (*Hippopotamus amphibius*) hair is sparsely present yet, sebaceous and sweat glands were so-far undetected (Figure 1) (Sokolov VE 1982; Graham 2005; Springer et al. 2021). In agreement, the collection of key molecular modules participating in sebum production in aquatic or semi-aquatic mammals was shown to mirror such mosaic of skin morphologies: being fully absent in cetaceans but only partially eroded in manatees and hippopotamuses (Lopes-Marques et al., 2019; Themudo et al., 2020; Springer et al. 2021). Extreme skin modifications are not restricted to aquatic species and include the naked mole rat and pangolins (Menon et al. 2019; Li et al. 2020; Savina et al. 2022). Species from the order Pholidota display a formidable keratin-scale armor probably serving as a key deterrent against predators and infections (Figure 1) (Meyer et al., 2013; Choo et al. 2016; Li et al., 2020). Interestingly, while pangolins have scattered hair on their abdomens, the presence of exocrine glands is contentious, since sweat and sebaceous glands were not detected in their dermis (Liumsiricharoen et al. 2008; Li et al. 2020). Consistently, the reported gene sequence decay of the melanocortin 5 receptor (*Mc5r*) in pangolins (Springer and Gatesy 2018; Liu et al. 2022), a gene with abundant expression in exocrine glands and centrally involved in sebogenesis (Eisinger et al. 2011; Xu et al. 2020; Shintani et al. 2021), is suggestive of a radical shift in skin exocrine function. Overall, the comparative molecular architecture governing skin physiology, in particular that associated with the glandular exocrine and eccrine compartment in Pholidota is largely unknown. Here, we provide an exhaustive comparative genomic analysis of the molecular modules governing mammalian skin homeostasis in four species of pangolins. Our findings provide a sharp insight into the richness of evolutionary routes and processes responsible for the skin physiology of extant mammalian lineages.

**Figure 1:**
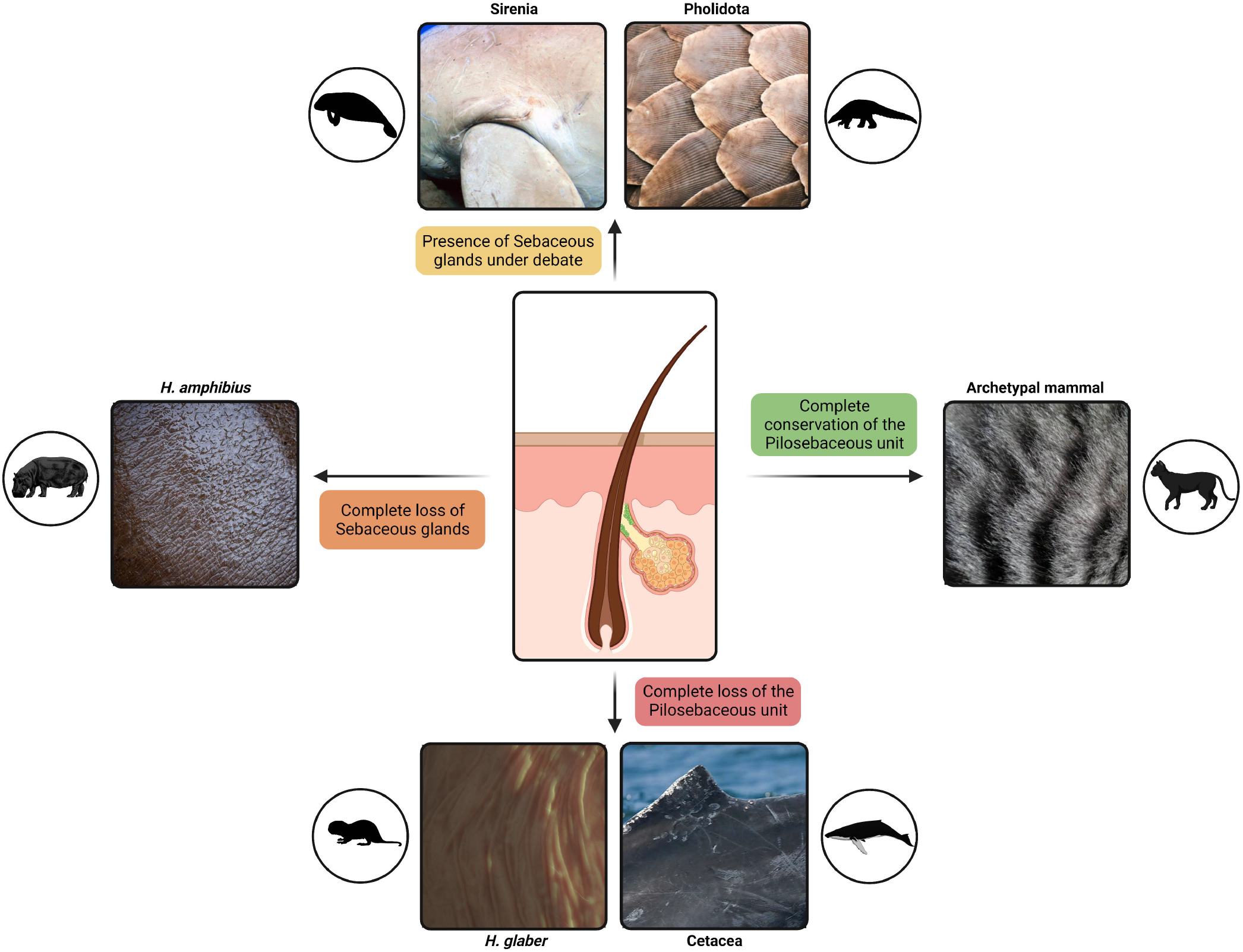
Schematic representation of the mammalian skin diversity, regarding the pilosebaceous gland. Cetaceans and the naked mole rat completely lack the pilosebaceous unit, while there is a presence of hair in hippopotamuses, although the sebaceous glands were also lost. In Sirenia and Pholidota, the presence and degree of functionality of this gland is disputed. Other mammalian species conserve the typical structure of the pilosebaceous gland. Created with BioRender.com

## Results and Discussion

In the present work, we set to analyze the status of skin-specific genes, mostly involved in sebaceous gland function, across four Pholidota species: *M. javanica, M. pentadactyla, M. crassicaudata* and *P. tricuspis*. Pangolins diversified approximately 38 million years ago and have since colonized two continents, Africa and Asia (Figure 2) (Gaubert et al. 2018). We first used PseudoIndex to classify the coding condition of the studied genes in pangolin genomes (Alves et al. 2020). This classification system varies in a discrete scale from 0 to 5, with 0 suggesting full functionality of the candidate gene and a value of 5 indicating a complete inactivation (Table 1). Genes with PseudoIndex above 2 were further validated and the retrieved mutations carefully annotated.

**Table 1:** PseudoIndex values (0 to 5, with 5 indicating probable mutations of pseudogenization) of the studied skin-related genes in 4 pangolin genomes. Genomic location per gene (Accession number) in each genome is indicated.

|  | *Manis crassicaudata*<br>GCA_016801295.1<br>N50 contig: 7,447 |  | *Manis javanica*<br>GCA_014570535.1<br>N50 contig: 83,964 |  | *Manis pentadactyla*<br>GCA_014570555.1<br>N50 contig: 151,937 |  | *Phataginus tricuspid*<br>GCA_004765945.1<br>N50 contig: 23,755 |  |
| --- | --- | --- | --- | --- | --- | --- | --- | --- |
| *AADACL3* | QZMM01052942.1 | X | NW_023436200.1 | 5 | NW_023456429.1 | 5 | SOZM010018664.1 | 5 |
| *AWAT1* | QZMM01191925.1 | X | NW_023436150.1 | 5 | NW_023454910.1 | 5 | SOZM010039185.1 | 5 |
| *AWAT2* | QZMM01012810.1 | 5 | NW_023436233.1 | 5 | NW_023457172.1 | 5 | SOZM010024681.1 | 5 |
| *DGAT2L6* | QZMM01083316.1<br>QZMM01012022.1<br>QZMM01201031.1 | 5 | NW_023436233.1 | 5 | NW_023454910.1 | 5 | SOZM010017370.1<br>SOZM010058764.1 | 5 |
| *EDA2R* | QZMM01332753.1<br>QZMM01003526.1<br>QZMM01124215.1<br>QZMM01281266.1 | X | NW_023436016.1 | 5 | NW_023458050.1 | 3 | SOZM010132249.1<br>SOZM010048560.1<br>SOZM010091528.1<br>SOZM010060741.1 | 5 |
| *FABP9* | QZMM01291322.1 | 5 | NW_023436188.1 | 5 | NW_023455931.1 | 5 | SOZM010004921.1 | 5 |
| *GSDMA* | QZMM01347780.1 | 5 | NW_023436081.1 | 5 | NW_023456908.1 | 0 | SOZM010000065.1 | 0 |
| *GSDMB* | QZMM01024159.1 | 5 | NW_023436081.1 | 5 | NW_023456908.1 | 5 | SOZM010000065.1 | 5 |
| *GSDMC* | QZMM01157587.1 | 5 | NW_023436052.1 | 1 | NW_023458035.1 | 5 | SOZM010244792.1 | 5 |
| *GSDMD* | QZMM01228481.1 | 3 | NW_023436087.1 | 0 | NW_023454130.1 | 0 | SOZM010008554.1 | 5 |
| *MOGAT3* | QZMM01311364.1 | 5 | NW_023436195.1 | 5 | NW_023454636.1 | 4 | SOZM010007253.1 | 5 |
| *SLURP1* | QZMM01029954.1 | 5 | NW_023436087.1 | X | NW_023454130.1 | X | SOZM010015272.1 | 3 |
| *TCHHL1* | QZMM01320808.1 | X | NW_023436233.1 | X | NW_023454669.1 | X | SOZM010020578.1<br>SOZM010004078.1 | X |

**Figure 2:**
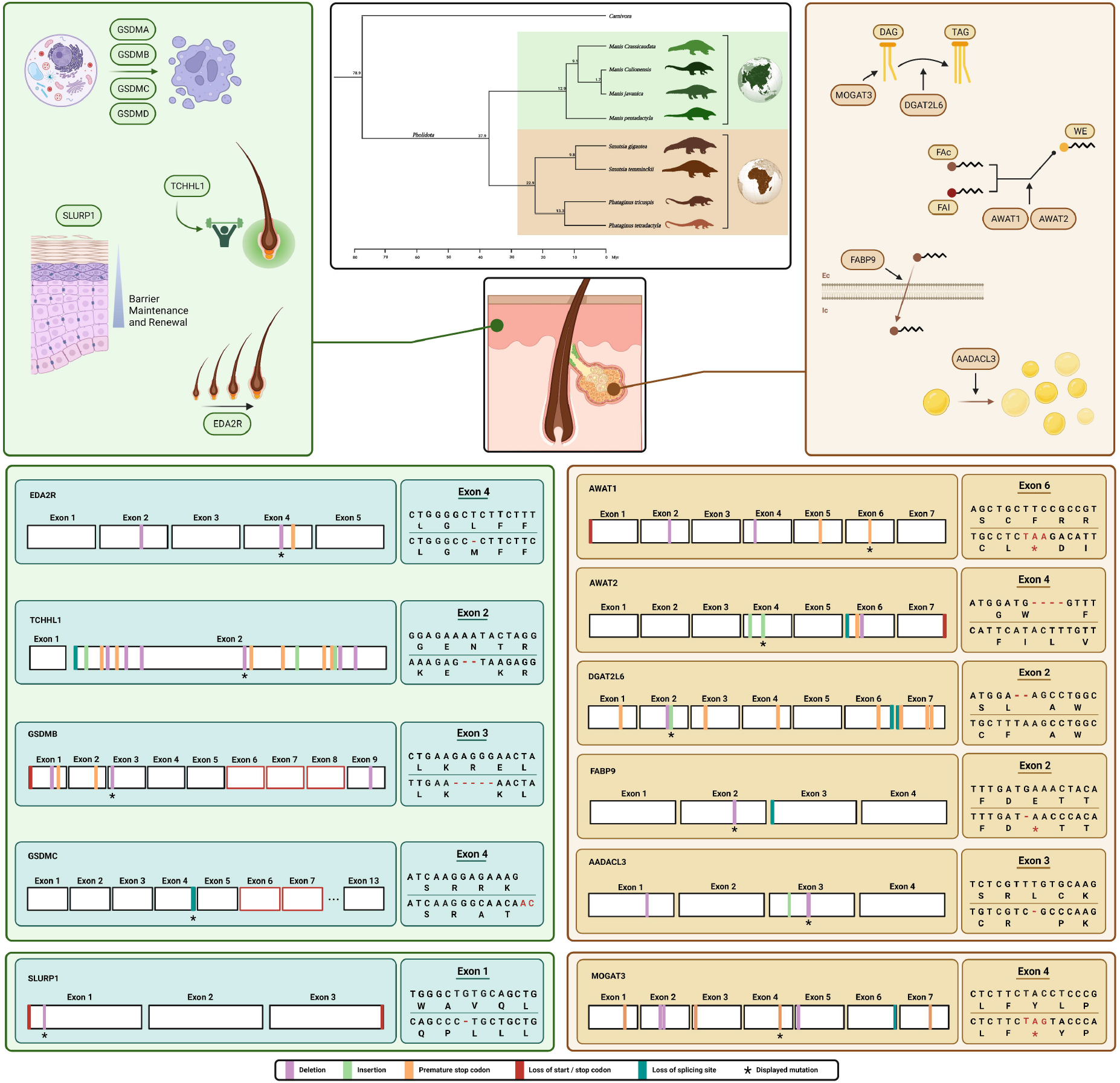
Mutational landscape of skin-genes in pangolins. Central top panel shows a phylogenetic tree of the evolution of Pholidota, and geographic distribution of extant species. On the upper half, in each side are represented the gene pathways: on the right side, genes associated with the production of sebum and lipid synthesis. On the left side, genes which play a role in skin defense, integrity of skin layers and homeostasis. On the lower half of the panel is described one conserved disruptive mutation for each gene, as well as a more global view of the numerous non-synonymous modifications along the gene. For *Slurp1* and *Mogat3*, no conserved mutations were detected, therefore the mutation displayed in the corresponding box is particular of a chosen species, mentioned alongside the illustration. Created with BioRender.com

### Sebum-producing genes show signs of erosion

Sebum comprises a complex lipid mixture. The biosynthesis of sebum components involves the action of various key modules: i.e., monoacylglycerol O-acyltransferases—*Mogat2* and *Mogat3*; diacyl-glycerol O-acyltransferases—*Dgat2* and *Dgat2l6*; wax alcohol acyltransferases—*Awat1* and *Awat2* (Bell and Coleman 1980; Turkish et al., 2005; Holmes 2010; Kawelke and Feussner 2015). Besides fatty acid esterification, to produce triglycerides or waxes, these modules also encompass fatty acid elongation (i.e. elongases—*Elovl3*), as well as trafficking and signaling (i.e. fatty acid binding protein—*Fabp9*), required for the upstream regulation of sebum production *via* fatty acid-responsive transcription factors (i.e. Peroxisome proliferator-activated receptors—PPARs) (Figure 2) (Trivedi et al. 2006; Kobayashi and Fujimori 2012).

Our comparative analysis showed that numerous disruptive mutations are present in the sebum-production related genes in pangolins. Regarding *Awat1*, we identified, and validated using independent SRA data, a conserved premature stop codon in the sixth exon in *M. javanica, M. pentadactyla* and *P. tricuspis* (Figure 2; Supplementary Material 1 and 2); in the *M. crassicaudata* genome no *Awat1* containing-scaffold was found yet, a possible problem in the genome assembly cannot be discarded (not shown). Other mutations, including nucleotide deletions and premature stop codons were also retrieved, notably a set of mutations conserved within the *Manis* genus (Supplementary material 1 and 2). Similarly to *Awat1, Awat2* displays a mutational pattern concurrent with the inactivation of the gene in the stem of the pangolin clade: exhibiting a conserved loss of a canonical splice site in exon 6 and lack of terminal stop codon in all examined species. Additionally, other disruptive mutations were mapped and validated: notably, a four-nucleotide insertion in the fourth exon found conserved within the *Manis* genus; or a 2-nucleotide deletion retrieved in the exon 4 of *P. tricuspis* (Figure 2; Supplementary Material 1 and 2). *Dgat2l6* orthologues also displayed several disruptive mutations in pangolins (Figure 2; Supplementary Material 1 and 2); yet, unlike *Awat1* and *Awat2*, none was shared across all pangolin species. Still, mutations were found to be conserved within the *Manis* genus (i.e. premature stops codons or the loss of splice sites in multiple different exons): notably a two-nucleotide insertion in the second exon, leading to a premature stop codon, which was further validated by SRA analysis (Figure 2; Supplementary Material 1 and 2). In *P. tricuspis*, a set of nucleotide deletions, premature stop codons, and a validated single-nucleotide insertion were identified (Supplementary Figure 1 and 2). Additionally, exon 1 of *P. tricuspis* and exon 5 of *M. pentadactyla* were not found (not shown). Such mutational patterns indicate an independent *Dgat2L6* erosion amongst pangolin lineages.

We next investigated *Fabp9* (Fatty Acid Binding Protein 9) and *Aadacl3* (Arylacetamide Deacetylase Like 3). *Fabp9* is typically expressed in the testis (Selvaraj et al. 2010), but previous findings have also suggested a role in skin homeostasis in Artiodactyls (Jiang et al. 2014). *Aadacl3*, although poorly studied, appears to be related with epidermal fat deposition (Lu et al. 2020; Sweet-Jones et al. 2021). Additionally, analysis of the Human Protein Atlas (www.proteinatlas.org) shows a very specific pattern with expression noted only in skin, breast, and placenta (not shown). Importantly, both these genes have been found to be inactivated on the stem Cetacea branch (Huelsmann et al. 2019; Monica Lopes-Marques et al. 2019; Springer et al. 2021). *Aadacl3* is also eroded in the African elephant (Huelsmann et al. 2019), a lineage where the presence of sebaceous glands has been contentious (Spearman 1970; Lopes-Marques et al. 2019). Sequence analysis allowed the identification of a conserved single nucleotide deletion in the third exon of *Aadacl3*, found in *M. javanica, M. pentadactyla* and *P. tricuspis*, leading to the emergence of a premature stop codon (Figure 2; Supplementary Material 1 and 2); as well as a conserved two nucleotide insertion in the same exon (Figure 2; Supplementary Material 1). No gene ORF was found for *M. crassicaudata* possibly due to low-quality genome assembly (not shown). Regarding *Fabp9*, a transversal canonical splice site loss was found in exon 3 for *M. javanica, M. pentadactyla* and *P. tricuspis*. Additional lineage-specific nucleotide deletions were also retrieved and validated (Figure 2; Supplementary Material 1 and 2). In *M. crassicaudata* the first exon was not found (Supplementary Figure 1). Finally, for *Mogat3*, no conserved mutation was detected, although the genomic sequences display several species-specific mutations across the four species: including, a premature stop codon on the sixth exon of *M. pentadactyla*, a single nucleotide deletion in the fourth exon of *M. crassicaudata* and numerous insertions in the second exon of *P. tricuspis* (Figure 2; Supplementary Material 1 and 2). During our analysis, we came across a possible case of gene duplication, followed by erosion of both copies of the gene (not shown), similarly to a previous duplication found in the hippopotamus (Springer et al. 2021). Altogether, or analyses show a comprehensive mutational landscape in sebum-related genes and highlight distinct evolutionary routes: with genes inactivated in the stem of the pangolin clade (i.e. *Awat1*), and genes convergently lost in both Asian and African lineages (i.e. *Dgat2l6*).

### *Elovl3* is functional and expressed in pangolin skin tissue compartments

Fatty acid elongation is a critical pathway for skin lipid homeostasis. A skin-specific elongase, *Elovl3*, has been shown to participate in the formation of specific and essential neutral lipids (Westerberg et al. 2004). In effect, the removal *Elovl3* in mice leads to a phenotype of sparse hair coat, hyperplastic pilosebaceous unit and perturbation of hair lipid contents (Westerberg et al. 2004). In Cetacea, *Elovl3* was previously shown to display inactivating mutations (Lopes-Marques et al. 2019; Springer et al. 2021). The examined pangolin *Elovl3* orthologues were all classified with a PseudoIndex of 0, an indication of sequence functionally (Table 1). Yet, we could not exclude deleterious mutations in the regulatory region of the gene that could hamper gene expression and function. Thus, we next examined the expression of *Elovl3* in a comprehensive panel of tissues (Figure 3; Supplementary Material 3). Of the examined tissues, the Pholidota *Elovl3* is uniquely expressed in the skin components, including hair follicles (Figure 3).

**Figure 3:**
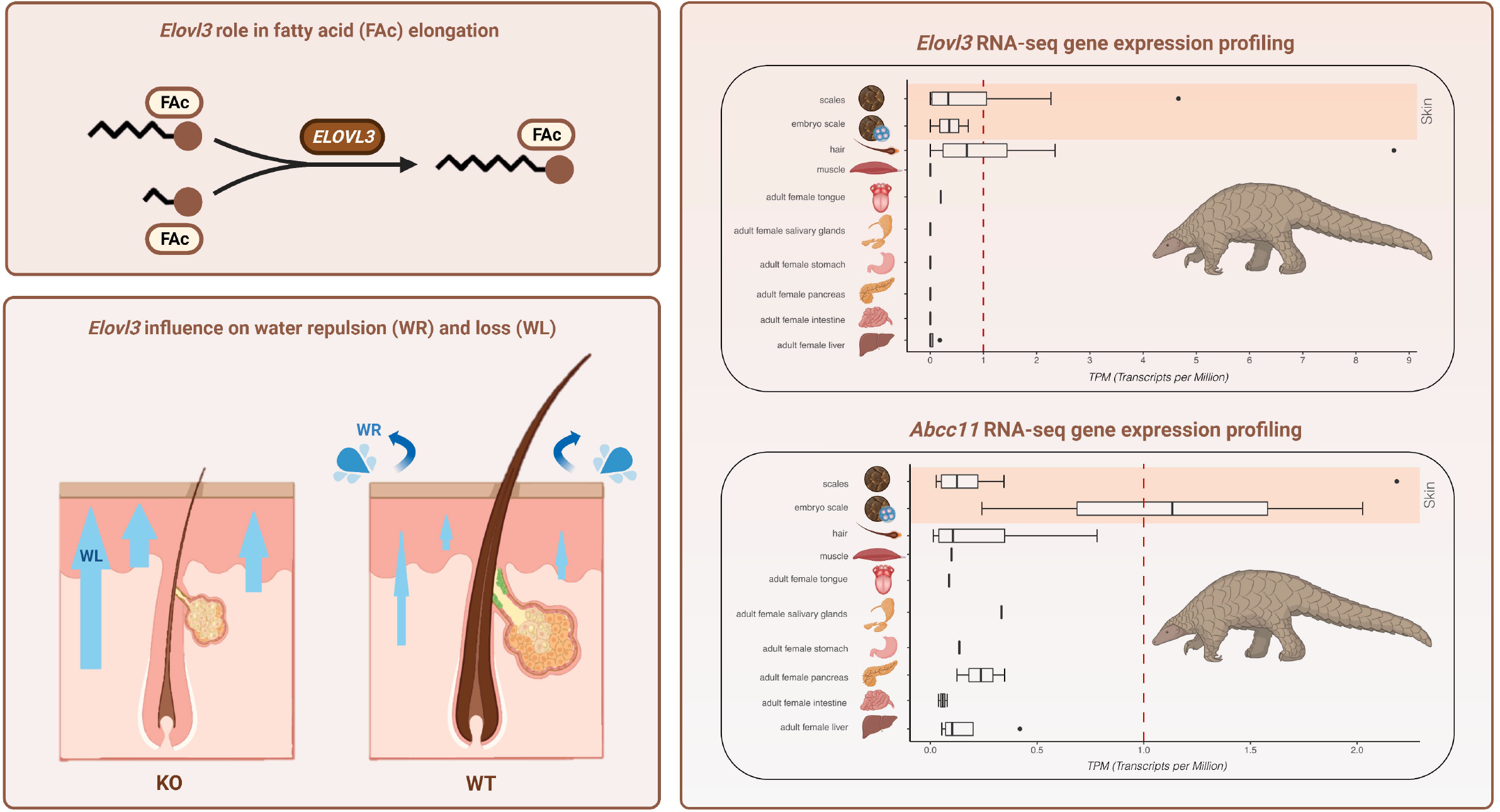
Tissue expression of *Elovl3* and *Abcc1* in the Java pangolin (M. *javanica*).

### Hair growth and robustness is shaped by gene loss episodes

We next expanded our analysis to a group of genes that pay a pivotal role in hair follicle homeostasis: Ectodysplasin A2 Receptor (*Eda2r*) and Trichohyalin Like 1 (*Tchhl1*). *Eda2r* is a membrane receptor which participates in the regulation of the hair follicle growth cycle (Kwacket al., 2019; Cai et al., 2021; Lan et al. 2020; Font-Porterias et al. 2022). Within the *Manis* genus, we detected a single nucleotide deletion in exon 2 and a premature stop codon in exon 4 of *Eda2r* (Figure 2; Supplementary Material 1 and 2); the latter was validated using independent SRAs (Supplementary Material 2). For *P. tricuspis*, a 2-nucleotide insertion in exon 3 and a premature stop codon in exon 5 were retrieved, denoting an independent *Eda2r* loss across Asian and African pangolin lineage. Additionally, multiple non-conserved alterations of the nucleotide sequence were identified in all four species: including insertions, deletions, premature stop codons, loss of splicing sites, and missing exons (Supplementary Material 1).

We next investigated *Tchhl1*, a gene responsible for providing mechanical strength to the hair follicle inner root sheath through keratin intermediate filaments (Makino et al. 2020). Numerous disruptive mutations were retrieved and found conserved amongst members of the *Manis* genus; a conserved two-nucleotide deletion in the second, and last exon of the gene, was selected for further scrutiny using independent SRAs (Figure 2; Supplementary Material 1 and 2). For *P. tricuspis*, no gene remnant was found. Curiously, *Tchhl1*, which is chiefly expressed in the *stratum basale* of the epidermal layer, is convergently inactivated in cetaceans and hippopotamuses (Springer et al. 2021).

### Erosion of the Gasdermin gene repertoire

Gasdermins comprise a protein family involved in membrane permeabilization and pyroptosis, a lytic pro-inflammatory type of cell death (Broz et al. 2019; de Schutter et al. 2021). Importantly, members of this gene family are expressed in the skin (Tamura et al. 2007). Our study of the gasdermin family (*Gsdma, Gsdmb, Gsdmc* and *Gsdmd*), revealed some very different scenarios. For *Gsdma* we only focused on the sequences of *M. crassicaudata* and *M. javanica*, as these scored over 2 in the preliminary PseudoIndex analysis, whereas *M. pentadactyla* and *P. tricuspis* scored 0, suggestive of intact genes (Table 1). Both species displayed the loss of one exon out of the eleven that constitute the gene, specifically the sixth exon (Supplementary Material 1). On the other hand, *Gsdmb* displayed various mutations, conserved across the *Manis* lineage, including a five-nucleotide deletion in the third exon, which was independently validated by SRAs (Supplementary Material 1 and 2). Other non-conserved mutations were identified and several exons could not be found through our analysis, especially in *P. tricuspis*, where only two out of the nine exons were retrieved, possibly due to a low-quality assembly (Figure 2). Yet, the *P. tricuspis* sequence also harbored a disruptive premature stop codon in exon 1, which was further validated (Supplementary Material 1 and 2). Regarding *Gsdmc*, no gene was found for *M. crassicaudata* and *P. tricuspis*; an assembly artifact cannot be discarded (Supplementary Material 4). Yet, for *M. javanica* and *M. pentadactyla* two *Gsdmc* copies were found, with all copies displaying Pseudoindex scores of 5 (Supplementary Material 4). Further analysis highlighted numerous inactivating mutations in all analyzed copies, confirming previous gene erosion predictions (Supplementary Material 4). For *Gsdmd*, Pseudoindex analysis on *M. javanica* and *Manis pentadactyla* yielded a value of 0, indicating a high level of conservation of the genomic sequence (Table 1). The evaluation of the remaining two species, on the other hand, revealed a case of exon loss in the sixth exon from a total of ten coding exons in *P. tricuspis*. In *M. crassicaudata*, the alignment similarity of the sixth exon was particularly low, hindering further conclusions (Supplementary Material 1 and 2).

### Skin-layer integrity genes display ORF-disruption mutations

Next, we investigated a gene related to the integrity of the skin layers—Secreted LY6/PLAUR Domain Containing 1 (*Slurp1*)—responsible for the stabilization of epithelial cell junctions (Campbell et al. 2019; Okamoto et al. 2020) and found to be eroded in Cetacea (Themudo et al. 2020). Mutations in this gene underlie a rare palmoplantar keratoderma exhibiting increased keratinocyte proliferation, lipid accumulation and water barrier deficiency (Fischer et al. 2001). We were unable to identify the *Slurp1*-containing scaffold in the *M. crassicaudata* genome (not shown). The remaining species showed solid evidence of pseudogenization: missing exons, in *M. javanica* and *M. pentadactyla*, loss of splicing sites in *M. pentadactyla*, or a deletion in the first exon of the gene in *P. tricuspis* (Figure 2). Validation of the mutations using independent SRAs was only possible for *P. tricuspis* (Supplementary Material 1 and 2).

### Sweat gland gene marker are functional in pangolins

Water evapotranspiration from the skin is fundamental for thermoregulation. This physiological process is dependent of the action of subset of skin elements, the sweat glands: eccrine, opening directly into the skin surface, and apocrine, opening into the pilosebaceous unit (Kobielak et al. 2015). Similarly to sebaceous glands, the presence of sweat glands in pangolins was so far unreported (Liumsiricharoen et al. 2008). Here, we investigated the expression status of *Abcc1*, a gene maker of apocrine sweat glands (Martin et al. 2010). *Abcc1* gene was previously shown to be eroded in Cetacea, paralleling sweat gland loss in this lineage (Oh et al. 2015). Expression analysis revealed a marked gene expression in the skin (Figure 3). Thus, despite the apparent absence of sweat glands, *Abcc1* was found intact in pangolins.

### Convergence, gene loss and the uniqueness of the skin phenotype in pangolins

Comparative genomics is a powerful tool to decipher the origin and loss of phenotypic variations (Huelsmann et al. 2019; Zoonomia Consortium 2020; Alves et al. 2021; Fuchs et al. 2022; Zheng et al. 2022). Specifically, gene loss-aware research is reverberating, highlighting the role of secondary losses in the emergence of diverse biological features: including the simplification of body plans (i.e., urochordates), the deconstruction of the vertebrate organs (i.e. stomach, pineal gland), the modulation of sensorial acuity (i.e. vision, taste), or even behavior and locomotion (Castro et al. 2013; Zhao et al. 2015; Lopes-Marques et al. 2019; Valente et al. 2021; Carneiro et al. 2021; Ferrández-Roldán et al. 2021; Indrischek et al. 2022).

Pholidota skin is unique among mammals. This group evolved from a common ancestral skin phenotype to armored keratinous scale appendices (Meyer et al. 2013). The main function of the keratinous scales is to serve as protection for the soft skin underneath, making it a natural shield against predators, and external harm such as UV radiation, as well as pathogenic agents (Wang et al. 2016). Additionally, the presence and functional status of sebaceous and sweat glands in Pholidota is so far unclear (Liumsiricharoen et al. 2008). Previous works emphasized gene-loss landscapes in other mammalian lineages with divergent skin phenotypes: such as cetaceans, and large African mammals such as the African elephant (*Loxodonta Africana*) or the white rhinoceros (*Ceratotherium sinum*) (Figure 4) (Plochocki et al. 2017; Springer and Gatesy 2018; Lopes-Marques et al. 2019; Springer et al., 2021); this prompted us to address whether similar genomic variations underlined the emergence of the distinctive skin phenotype in pangolins. Our comparative analysis led to the annotation and validation of numerous disruptive mutations in target genes—related with sebum production, skin layer development and maintenance, and hair growth—across four studied species spanning the two pangolin lineages. These results support the role of gene pseudogenization episodes as drivers of the extant skin phenotype in Pholidota. Ancestral (i.e. *Awat1, Awat2, Aacadl3*) and convergent (i.e. *Dgat2l6*) pseudogenization events in genes associated with epidermal lipids and sebum production strongly suggest a progressive impairment of the sebum producing molecular machinery in pangolin lineages. Similar genomic signatures were previously proposed for mammalian lineages with derived skin phenotypes, particularly visible in the fully-aquatic Cetacea (Sokolov VE 1982; Springer and Gatesy 2018; Lopes-Marques et al. 2019). Sebum, a mammalian synapomorphy, is mainly produced to serve as a protective layer against UV radiation, bacteria, and skin dehydration (Lobitz 1957; Pappas 2009; Niemann and Horsley 2012). These functions, although vital when considering exposed skin, may have diminished relevance in an armor-like skin, as observed in Pholidota (Figure 4). A similar hypothesis can be drawn regarding genes related to skin protection against external dangers (i.e. microbial infection)—the gasdermin family, responsible for host defense and cell death (Tamura and Shiroishi 2015).

**Figure 4:**
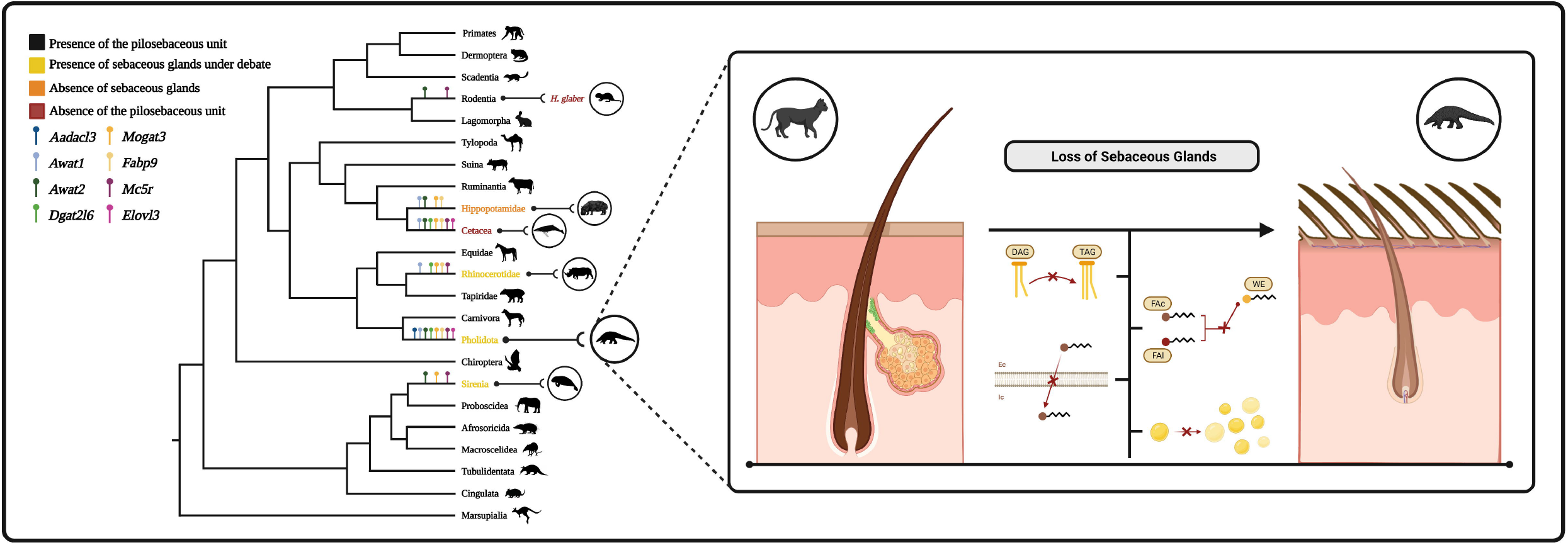
Schematic illustration of the loss of sebaceous glands in mammals and associated gene loss events. Phylogenetic tree of mammals, with emphasis to the different sebaceous glands-related phenotypes. In each branch of the tree are highlighted the different genes each mammalian group lost during the course of its evolutionary development. Each different skin phenotype is characterized by a particular color. Each gene is symbolized with a particular coloration as well. Created with BioRender.com

Whereas sebaceous gland dismantling is supported by our data, the current results do not unequivocally clarify the fate of sweat glands in pangolins. *Abcc1* gene, a marker of apocrine sweat glands (Martin et al. 2010), generally associated with the hair follicle, was shown to be intact and expressed in pangolins. Yet, unlike their back and tail, the abdomen of pangolins yields an exposed skin with sparse hair and thicker *stratum corneum* (Meyer et al. 2013). In agreement, evidence of erosion was found in genes related with keratinocyte proliferation and stabilization (i.e. *Slurp1*) or hair development and mechanical strength (i.e. *Edar2, Tchhl1*) (Campbell et al. 2019; Lan et al. 2020; Cai et al. 2021). Conversely, other genomic components were found intact. Among these we find *Alox15A* and *Alox3*, epidermal lipoxygenases participating in the maintenance of the cornified layer (Krieg et al. 2013) (not shown), or *Elovl3*. Such mosaic gene retention has been previously proposed for mammalian species with derived skin phenotypes (i.e. elephant, rhinoceros, hippopotamuses) (Lopes-Marques et al. 2019; Springer et al. 2021).

In conclusion, our findings show that species of the order Pholidota display numerous skin-related gene pseudogenization events paralleled by the dismantling of the sebaceous gland in this mammalian group and the emergence of their idiosyncratic skin phenotype. Importantly, the present work reinforces the role of gene loss as a powerful evolutionary driver, notably in transitional scenarios or radical phenotypic adaptation, as reported for other mammalian groups such as the fully aquatic Cetacea and Sirenia.

## Material and Methods

### Genome Resources

The genome regions of the target genes were retrieved from the NCBI (National Center for Biotechnology Information) database. Two of the studied species have high-quality genome annotations (*M. javanica* (GCA_014570535.1) and *M. pentadactyla* (GCA_014570555.1) (Table 1). Thus, target genes and corresponding genomic regions were retrieved using gene symbols and manually verified. For *M. crassicaudata* (GCA_016801295.1) and *P. tricuspis* (GCA_004765945.1) (Table 1), given that no genome annotation is available, we used BLAST (Basic Local Alignment Search Tool), using as query the target and two flanking genes from species with annotated genomes, to extract the full scaffold sequence. Gene selection involved an exhaustive literature review to define key genetic pathways involved in skin physiology (e.g. Themudo et al., 2020). This initial screening was complemented with the investigation of the skin “*enriched genes*” list from the Human Protein Atlas resource (Uhlén et al. 2015). Additionally, we used *String* to explore protein-protein networks of skin-specific gene families (Szklarczyk et al. 2021).

### Sequence Alignment Analysis

Each gene sequence was aligned against a reference coding and curated sequence for each gene (*Homo sapiens*) using PseudoChecker (Alves et al. 2020), a program that uses the MACSE alignment software (Ranwez et al. 2018). PseudoChecker attributes a value ranging from 0 to 5, the PseudoIndex, with respect to the status of the genomic sequence in comparison to the reference: a value of 0 representing a fully functional protein coding sequence and a value of 5 suggesting pseudogenization mutations (Table 1). For the genomic regions scoring 3 or higher on PseudoIndex, a second alignment was performed, against the same reference, using Geneious Prime 2021.2.2 (https://www.geneious.com), for manual curation and evaluation.

### Mutational Validation *via* SRAs

Identified mutations were validated using Sequence Read Archives (SRAs) (Supplementary material 5). Two independent projects were used *per* species when possible and aligned with our genomic sequence using Geneious Prime 2021.2.2. Exceptions include *P. tricuspis*, for which only one SRA is available and *M. crassicaudata* for which no SRA project exist. Cross-species conserved non-synonymous mutations were selected for further validation and, when conservation was authenticated, mutations located in the mid-section of the gene sequence were selected, due to the increased stability of gene structure and lower susceptibility to alignment artifacts.

### Gene Expression analysis of *Elovl3* and *Abcc11* in pangolin tissues

RNA-seq data for the tissues of *M. javanica* from previously published bioprojects were downloaded from NCBI’s database (Supplementary Material 3). In order to map the SRAs to the reference genome, we indexed the genome using Hisat2 v.2.2.0 (Kim et al. 2015; Kim et al. 2019; Zhang et al. 2021) to enable the mapping of both forward and reverse sequences (within the SRA archives) to the species’ reference genome (YNU_ManJav_2.0). In this mapping, the option for downstream transcriptome assembly was triggered to reduce computational and memory consumption with transcript assembly. The generated SAM files were then converted and sorted to BAM files using samtools and submitted to featureCounts, to quantify the raw reads that were mapped to the transcriptomic components (Liao et al. 2014); the raw read quantification was then transformed to transcripts per million (TPM) using an in-house script.

## References

Alves LQ, Ruivo R, Fonseca MM, Lopes-Marques MO, Ribeiro P, Castro LFC. 2020. PseudoChecker: An integrated online platform for gene inactivation inference. Nucleic Acids Res. 48:321–331.

Alves LQ, Ruivo R, Valente R, Fonseca MM, Machado AM, Plön S, Monteiro N, García-Parraga D, Ruiz-Díaz S, Sánchez-Calabuig Mj, et al. 2021. A drastic shift in the energetic landscape of toothed whale sperm cells. Current Biology 31:3648–3655.

Bell RM, Coleman RA. 1980. ENZYMES OF GLYCEROLIPID SYNTHESIS IN EUKARYOTES. Annu Rev Biochem 49:459–87

Broz P, Pelegrín P, Shao F. 2019. The gasdermins, a protein family executing cell death and inflammation. Nat Rev Immunol. 2019 20:3

Cai Z, Deng X, Jia J, Wang D, Yuan G. 2021. Ectodysplasin A/Ectodysplasin A Receptor System and Their Roles in Multiple Diseases. Front Physiol. 12.

Campbell G, Swamynathan S, Tiwari A, Swamynathan SK. 2019. The secreted Ly-6/uPAR related protein-1 (SLURP1) stabilizes epithelial cell junctions and suppresses TNF-α-induced cytokine production. Biochem Biophys Res Commun 517:729–734.

Carneiro M, Vieillard J, Andrade P, Boucher S, Afonso S, Blanco-Aguiar JA, Santos N, Branco J, Esteves PJ, Ferrand N, et al. 2021. A loss-of-function mutation in RORB disrupts saltatorial locomotion in rabbits. PLoS Genetics.

Castro LFC, Gonçalves O, Mazan S, Tay BH, Venkatesh B, Wilson JM. 2013. Recurrent gene loss correlates with the evolution of stomach phenotypes in gnathostome history. Proc. R. Soc. B. 281.

Choo SW, Rayko M, Tan TK, Hari R, Komissarov A, Wee WY, Yurchenko AA, Kliver S, Tamazian G, Antunes A, et al. 2016. Pangolin genomes and the evolution of mammalian scales and immunity. Genome Res. 26:1312–1322.

Eisinger M, Li WH, Anthonavage M, Pappas A, Zhang L, Rossetti D, Huang QL, Seiberg M. 2011. A melanocortin receptor 1 and 5 antagonist inhibits sebaceous gland differentiation and the production of sebum-specific lipids. J Dermatol Sci. 63:23– 32.

Ferrández-Roldán A, Fabregà-Torrus M, Sánchez-Serna G, Duran-Bello E, Joaquín-Lluís M, Bujosa P, Plana-Carmona M, Garcia-Fernàndez J, Albalat R, Cañestro C. 2021. Cardiopharyngeal deconstruction and ancestral tunicate sessility. Nature 599:431– 435.

Fischer J, Bouadjar B, Heilig R, Huber M, Lefèvre C, Jobard F, Macari F, Bakija-Konsuo A, Ait-Belkacem F, Weissenbach J, et al. 2001. Mutations in the gene encoding SLURP-1 in Mal de Meleda. Hum Mol Genet. 10:875–880.

Font-Porterias N, McNelis MG, Comas D, Hlusko LJ. 2022. Evidence of selection in the ectodysplasin pathway among endangered aquatic mammals. Integrative Organismal Biology.

Fuchs P, Drexler C, Ratajczyk S, Eckhart L. 2022. Comparative genomics reveals evolutionary loss of epiplakin in cetaceans. Scientific Reports 12:1–9.

Gaubert P, Antunes A, Meng H, Miao L, Peigné S, Justy F, Njiokou F, Dufour S, Danquah E, Alahakoon J, et al. 2018. The Complete Phylogeny of Pangolins: Scaling Up Resources for the Molecular Tracing of the Most Trafficked Mammals on Earth. J Hered. 109:347–359.

Graham A-R. 2005. HISTOLOGICAL EXAMINATION OF THE FLORIDA MANATEE (Trichecus manatus latirostris) INTEGUMENT. Gainesville, Florida

Holmes RS. 2010. Comparative genomics and proteomics of vertebrate diacylglycerol acyltransferase (DGAT), acyl CoA wax alcohol acyltransferase (AWAT) and monoacylglycerol acyltransferase (MGAT). Comp Biochem Physiol Part D Genomics Proteomics 5:45–54.

Holthaus KB, Lachner J, Ebner B, Tschachler E, Eckhart L. 2021. Gene duplications and gene loss in the epidermal differentiation complex during the evolutionary land-to-water transition of cetaceans. Scientific Reports 11:1–9.

Huelsmann M, Hecker N, Springer MS, Gatesy J, Sharma V, Hiller M. 2019. Genes lost during the transition from land to water in cetaceans highlight genomic changes associated with aquatic adaptations. Sci Adv. 5.

Indrischek H, Hammer J, Machate A, Hecker N, Kirilenko B, Roscito J, Hans S, Norden C, Brand M, Hiller M. 2022. Vision-related convergent gene losses reveal SERPINE3’s unknown role in the eye. Elife 11.

Jiang Y, Xie M, Chen W, Talbot R, Maddox JF, Faraut T, Wu C, Muzny DM, Li Y, Zhang W, et al. 2014. The sheep genome illuminates biology of the rumen and lipid metabolism. Science 344:1168–1173.

Kawelke S, Feussner I. 2015. Two predicted transmembrane domains exclude very long chain fatty acyl-CoAs from the active site of mouse wax synthase. PLoS One 10.

Kim D, Langmead B, Salzberg SL. 2015. HISAT: a fast spliced aligner with low memory requirements. Nat Methods 12:357–360.

Kim D, Paggi JM, Park C, Bennett C, Salzberg SL. 2019. Graph-based genome alignment and genotyping with HISAT2 and HISAT-genotype. Nature Biotechnology 37:907–915.

Kobayashi T, Fujimori K. 2012. Very long-chain-fatty acids enhance adipogenesis through coregulation of Elovl3 and PPARγ in 3T3-L1 cells. Am J Physiol Endocrinol Metab. 302.

Kobielak K, Kandyba E, Leung Y. 2015. Skin and Skin Appendage Regeneration. In: Translational Regenerative Medicine. Elsevier. p. 269–292.

Kowalczyk A, Chikina M, Clark N. 2021. Complementary evolution of coding and noncoding sequence underlies mammalian hairlessness. bioRxiv.

Krieg P, Rosenberger S, de Juanes S, Latzko S, Hou J, Dick A, Kloz U, van der Hoeven F, Hausser I, Esposito I, et al. 2013. Aloxe3 knockout mice reveal a function of epidermal lipoxygenase-3 as hepoxilin synthase and its pivotal role in barrier formation. J Invest Dermatol. 133:172–180.

Lan X, Kumar V, Jha A, Aslam R, Wang H, Chen K, Yu Y, He W, Chen F, Luo H, et al. 2020. EDA2R mediates podocyte injury in high glucose milieu. Biochimie 174:74– 83.

Li HM, Liu P, Zhang XJ, Li LM, Jiang HY, Yan H, Hou FH, Chen JP. 2020. Combined proteomics and transcriptomics reveal the genetic basis underlying the differentiation of skin appendages and immunity in pangolin. Scientific Reports 10:1–13.

Liao Y, Smyth GK, Shi W. 2014. featureCounts: an efficient general purpose program for assigning sequence reads to genomic features. Bioinformatics 30:923–930.

Liu J, Shu M, Liu S, Xue J, Chen H, Li W, Zhou J, Amanullah A, Guan M, Bao J, et al. 2022. Differential MC5R loss in whales and manatees reveals convergent evolution to the marine environment. Dev Genes Evol. 232:81–87.

Liumsiricharoen M, Prapong T, Chungsamarnyart N, Thiangtum K, Pongket P, Ruengsuphaphichat P, Suprasert A. 2008. Macroscopic and Microscopic Study of the Integument and Accessory Organs of Malayan Pangolin (Manis javanica). Kasetsart Veterinarians.

Lobitz WC. 1957. The Structure and Function of the Sebaceous Glands. AMA Arch Derm.

Lopes-Marques Monica, Machado AM, Alves LQ, Fonseca MM, Barbosa S, Sinding MHS, Rasmussen MH, Iversen MR, Bertelsen MF, Campos PF, et al. 2019. Complete inactivation of sebum-producing genes parallels the loss of sebaceous glands in Cetacea. Mol Biol Evol. 36:1270–1280.

Lopes-Marques Mónica, Ruivo R, Alves LQ, Sousa N, Machado AM, Castro LFC. 2019. The singularity of cetacea behavior parallels the complete inactivation of melatonin gene modules. Genes (Basel) 10.

Lu Z, Yue Y, Yuan C, Liu J, Chen Z, Niu C, Sun X, Zhu S, Zhao H, Guo T, et al. 2020. Genome-Wide Association Study of Body Weight Traits in Chinese Fine-Wool Sheep. Animals 10.

Makino T, Mizawa M, Yoshihisa Y, Yamamoto S, Tabuchi Y, Miyai M, Hibino T, Sasahara M, Shimizu T. 2020. Trichohyalin-like 1 protein plays a crucial role in proliferation and anti-apoptosis of normal human keratinocytes and squamous cell carcinoma cells. Cell Death Discov. 6.

Martin A, Saathoff M, Kuhn F, Max H, Terstegen L, Natsch A. 2010. A Functional ABCC11 Allele Is Essential in the Biochemical Formation of Human Axillary Odor. JID 130:529–540.

Menon GK, Catania KC, Crumrine D, Bradley C, Mauldin EA. 2019. Unique features of the skin barrier in naked mole rats reflect adaptations to their fossorial habitat. J Morphol. 280:1871–1880.

Meyer W, Liumsiricharoen M, Suprasert A, Fleischer LG, Hewicker-Trautwein M. 2013. Immunohistochemical demonstration of keratins in the epidermal layers of the Malayan pangolin (Manis javanica), with remarks on the evolution of the integumental scale armour. Eur J Histochem. 57:172–177.

Nery MF, Arroyo JI, Opazo JC. 2014. Increased rate of hair keratin gene loss in the cetacean lineage. BMC Genomics 15:1–9.

Niemann C, Horsley V. 2012. Development and homeostasis of the sebaceous gland. Semin Cell Dev Biol. 23:928–936.

Oh JW, Chung O, Cho YS, Macgregor GR, Plikus M v. 2015. Gene loss in keratinization programs accompanies adaptation of Cetacean skin to aquatic lifestyle. Exp Dermatol. 24:572.

Okamoto R, Goto I, Nishimura Y, Kobayashi I, Hashizume R, Yoshida Y, Ito R, Kobayashi Y, Nishikawa M, Ali Y, et al. 2020. Gap junction protein beta 4 plays an important role in cardiac function in humans, rodents, and zebrafish. PLoS One 15.

Pappas A. 2009. Epidermal surface lipids. Dermatoendocrinol 1:72–76.

Plochocki JH, Ruiz S, Rodriguez-Sosa JR, Hall MI. 2017. Histological study of white rhinoceros integument. PLoS One 12.

Ranwez V, Douzery EJP, Cambon C, Chantret N, Delsuc F. 2018. MACSE v2: Toolkit for the alignment of coding sequences accounting for frameshifts and stop codons. Mol Biol Evol. 35:2582–2584.

Reeb D, Best PB, Kidson SH. 2007. Structure of the integument of southern right whales,Eubalaena australis. The Anatomical Record: Advances in Integrative Anatomy and Evolutionary Biology 290:596–613.

Savina A, Jaffredo T, Saldmann F, Faulkes CG, Moguelet P, Leroy C, Marmol D del, Codogno P, Foucher L, Zalc A, et al. 2022. Single-cell transcriptomics reveals age-resistant maintenance of cell identities, stem cell compartments and differentiation trajectories in long-lived naked mole-rats skin. Aging 14:3728–3756.

de Schutter E, Roelandt R, Riquet FB, van Camp G, Wullaert A, Vandenabeele P. 2021. Punching Holes in Cellular Membranes: Biology and Evolution of Gasdermins. Trends Cell Biol. 31:500–513.

Selvaraj V, Asano A, Page JL, Nelson JL, Kothapalli KSD, Foster JA, Brenna JT, Weiss RS, Travis AJ. 2010. Mice lacking FABP9/PERF15 develop sperm head abnormalities but are fertile. Dev Biol. 348:177.

Shintani A, Sakata-Haga H, Moriguchi K, Tomosugi M, Sakai D, Tsukada T, Taniguchi M, Asano M, Shimada H, Otani H, et al. 2021. MC5R Contributes to Sensitivity to UVB Waves and Barrier Function in Mouse Epidermis. JID Innov. 1:100024.

Sokolov VE. 1982. Mammal Skin. London (CA): University of California Press

Spearman RI. 1970. The Epidermis and its Keratinisation in the African Elephant (Loxodonta Africana). Zoologica Africana 5:327–338.

Spearman RI. 1972. The epidermal stratum corneum of the whale. J Anat 113:373– 381.

Springer MS, Gatesy J. 2018. Evolution of the MC5R gene in placental mammals with evidence for its inactivation in multiple lineages that lack sebaceous glands. Mol Phylogenet Evol. 120:364–374.

Springer MS, Guerrero-Juarez CF, Huelsmann M, Collin MA, Danil K, McGowen MR, Oh JW, Ramos R, Hiller M, Plikus M v., et al. 2021. Genomic and anatomical comparisons of skin support independent adaptation to life in water by cetaceans and hippos. Current Biology 31:2124–2139.

Sweet-Jones J, Yurchenko AA, Igoshin A v., Yudin NS, Swain MT, Larkin DM. 2021. Resequencing and signatures of selection scan in two Siberian native sheep breeds point to candidate genetic variants for adaptation and economically important traits. Anim Genet. 52:126–131.

Szklarczyk D, Gable AL, Nastou KC, Lyon D, Kirsch R, Pyysalo S, Doncheva NT, Legeay M, Fang T, Bork P, et al. 2021. The STRING database in 2021: customizable protein– protein networks, and functional characterization of user-uploaded gene/measurement sets. Nucleic Acids Res. 49:605.

Tamura M, Shiroishi T. 2015. GSDM family genes meet autophagy. Biochemical Journal 469:5–7.

Tamura M, Tanaka S, Fujii T, Aoki A, Komiyama H, Ezawa K, Sumiyama K, Sagai T, Shiroishi T. 2007. Members of a novel gene family, Gsdm, are expressed exclusively in the epithelium of the skin and gastrointestinal tract in a highly tissue-specific manner. Genomics 89:618–629.

Themudo G, Alves LQ, Machado AM, Lopes-Marques M, da Fonseca RR, Fonseca M, Ruivo R, Castro LFC. 2020. Losing Genes: The Evolutionary Remodeling of Cetacea Skin. Front Mar Sci. 7.

Trivedi NR, Cong Z, Nelson AM, Albert AJ, Rosamilia LL, Sivarajah S, Gilliland KL, Liu W, Mauger DT, Gabbay RA, et al. 2006. Peroxisome proliferator-activated receptors increase human sebum production. J Invest Dermatol. 126:2002–2009.

Uhlén M, Fagerberg L, Hallström BM, Lindskog C, Oksvold P, Mardinoglu A, Sivertsson Å, Kampf C, Sjöstedt E, Asplund A, et al. 2015. Tissue-based map of the human proteome. Science 347.

Valente R, Alves F, Sousa-Pinto I, Ruivo R, Castro LFC. 2021. Functional or Vestigial? The Genomics of the Pineal Gland in Xenarthra. J Mol Evol. 89:565–575.

Valente R, Alves LQ, Nabais M, Alves F, Sousa-Pinto I, Ruivo R, Castro LFC. 2021. Convergent Cortistatin losses parallel modifications in circadian rhythmicity and energy homeostasis in Cetacea and other mammalian lineages. Genomics 113:1064– 1070.

Wang B, Yang W, Sherman VR, Meyers MA. 2016. Pangolin armor: Overlapping, structure, and mechanical properties of the keratinous scales. Acta Biomater 41:60– 74.

Westerberg R, Tvrdik P, Undén AB, Månsson JE, Norlén L, Jakobsson A, Holleran WH, Elias PM, Asadi A, Flodby P, et al. 2004. Role for ELOVL3 and fatty acid chain length in development of hair and skin function. J Biol Chem. 279:5621–5629.

Wu T, Deme L, Zhang Z, Huang X, Xu S, Yang G. 2022. Decay of TRPV3 as the genomic trace of epidermal structure changes in the land-to-sea transition of mammals. Ecol Evol 12.

Xu Y, Guan X, Zhou R, Gong R. 2020. Melanocortin 5 receptor signaling pathway in health and disease. Cell. Mol. Life Sci.77:3831–3840.

Zhang Y, Park C, Bennett C, Thornton M, Kim D. 2021. Rapid and accurate alignment of nucleotide conversion sequencing reads with HISAT-3N. Genome Res. 31:1290– 1295.

Zhao H, Li J, Zhang J. 2015. Molecular evidence for the loss of three basic tastes in penguins. Current Biology 25:141–142.

Zheng Z, Hua R, Xu G, Yang H, Shi P. 2022. Gene losses may contribute to subterranean adaptations in naked mole-rat and blind mole-rat. BMC Biol. 20:1–17.

Zoonomia Consortium. 2020. A comparative genomics multitool for scientific discovery and conservation. Nature 587:240–245.

