## Supplemental Material 2 for "Convergent Decay of Skin-specific Gene Modules in Pangolins"

SRA Validation of an insertion in exon 3 of *Aadacl3*:

*Manis javanica:*

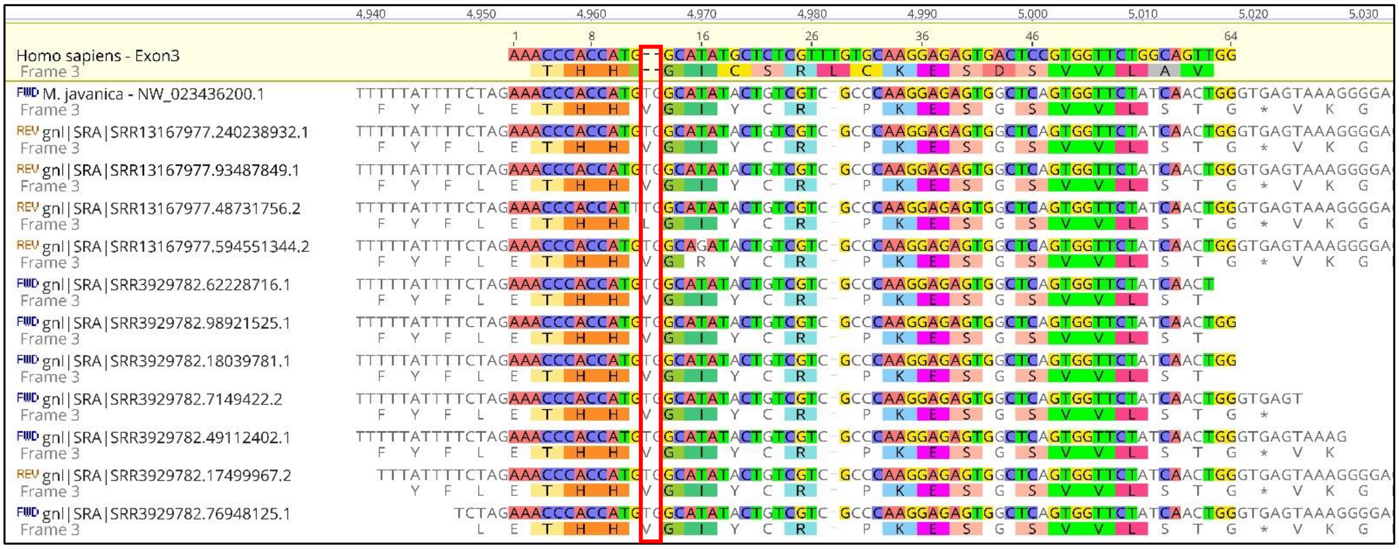


*Manis pentadactyla:*

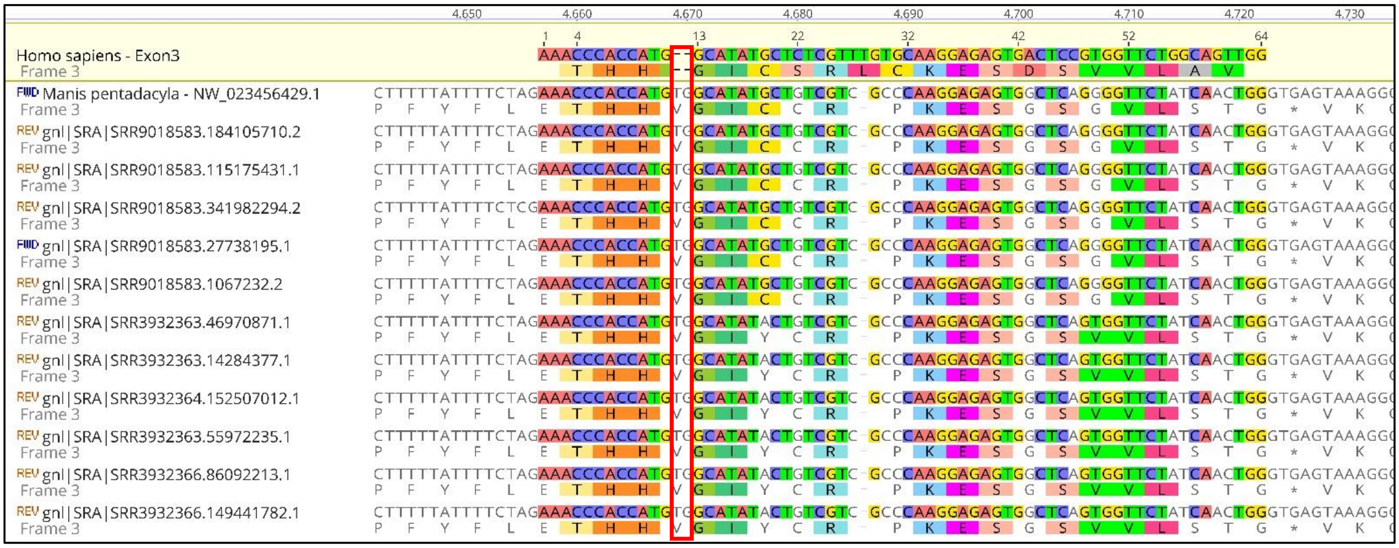

*Phataginus tricuspis:*

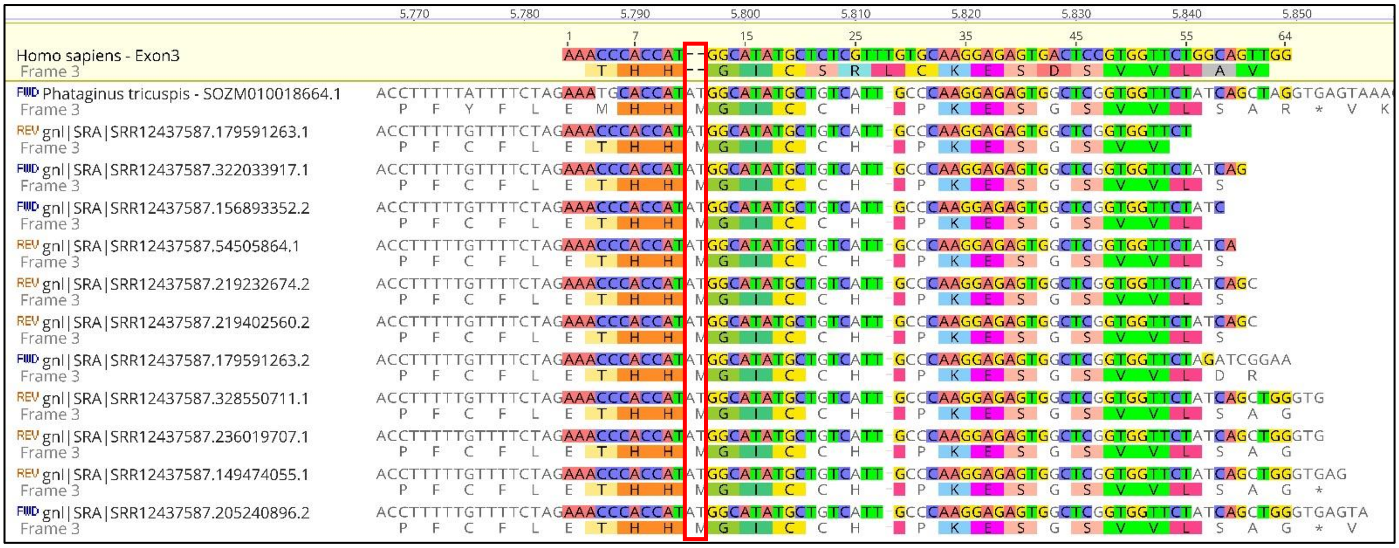


SRA Validation of a premature stop codon in exon 6 of *Awat1*:

*Manis javanica:*

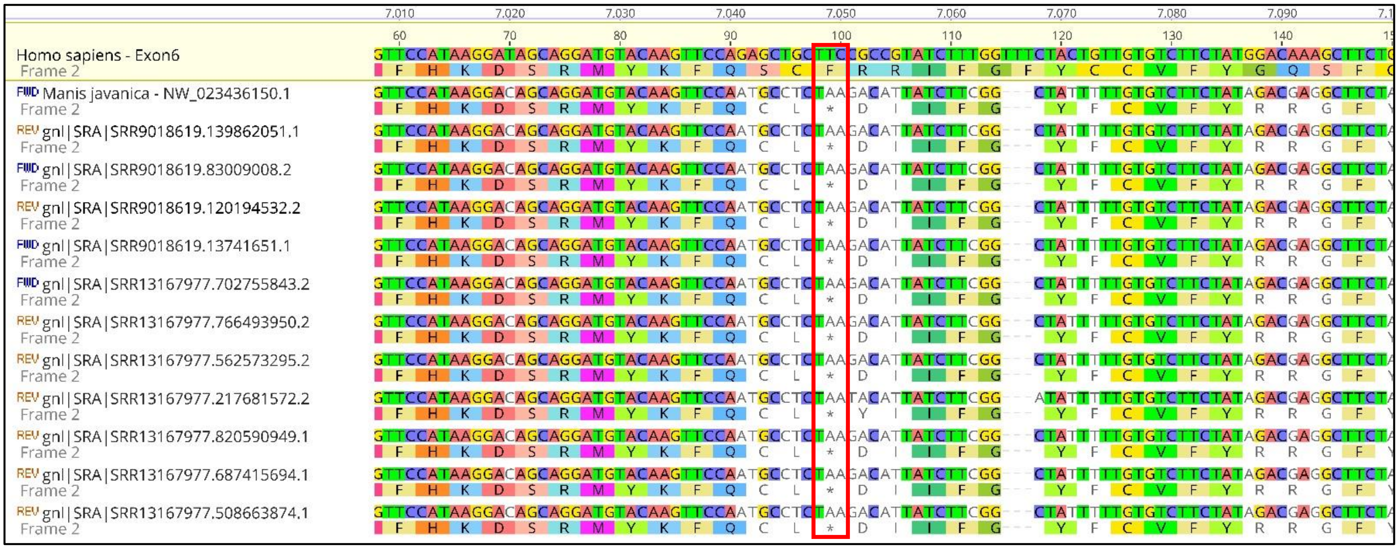

*Manis pentadactyla:*

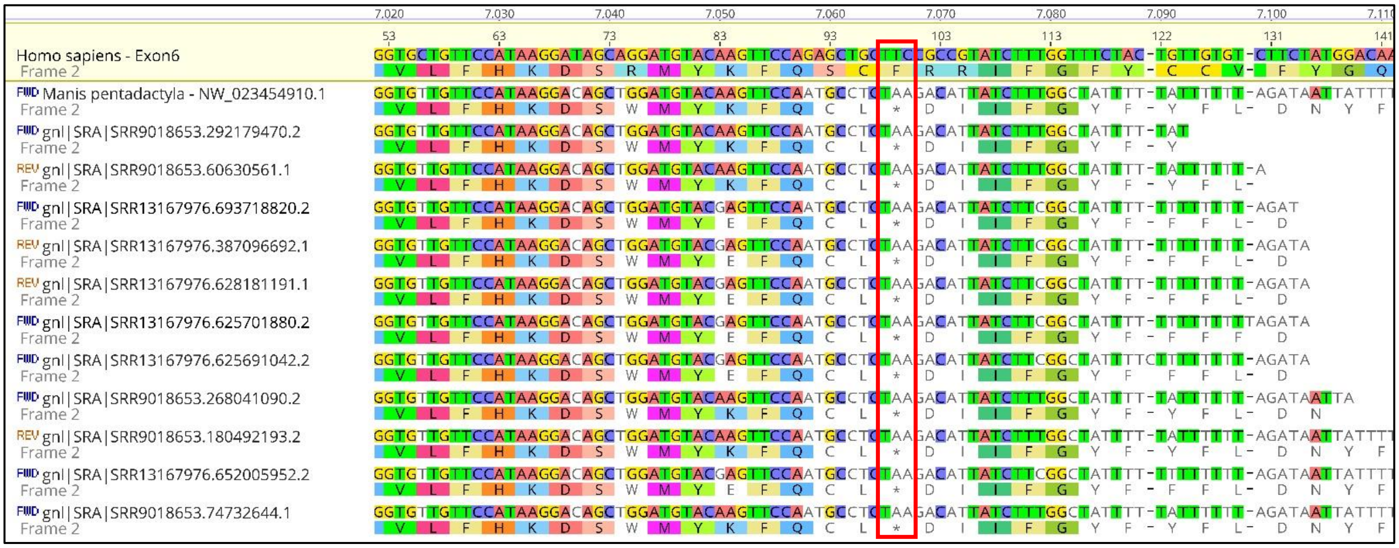

*Phataginus tricuspis:*

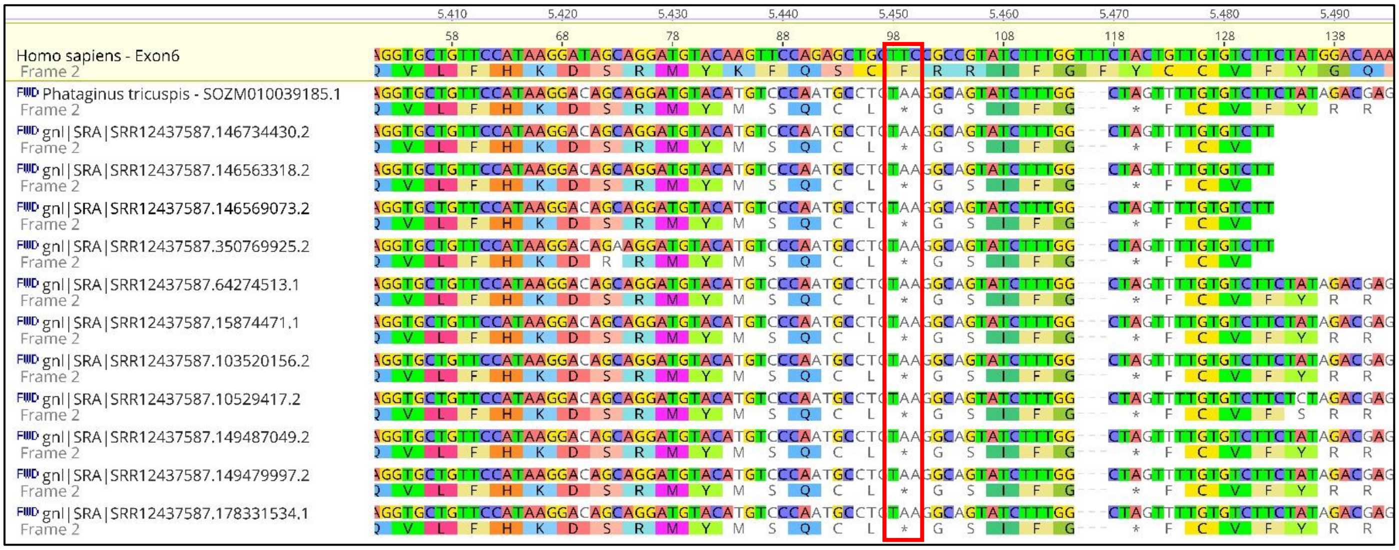

SRA Validation of an insertion in exon 4 of *Awat2* in Manis:

*Manis javanica:*

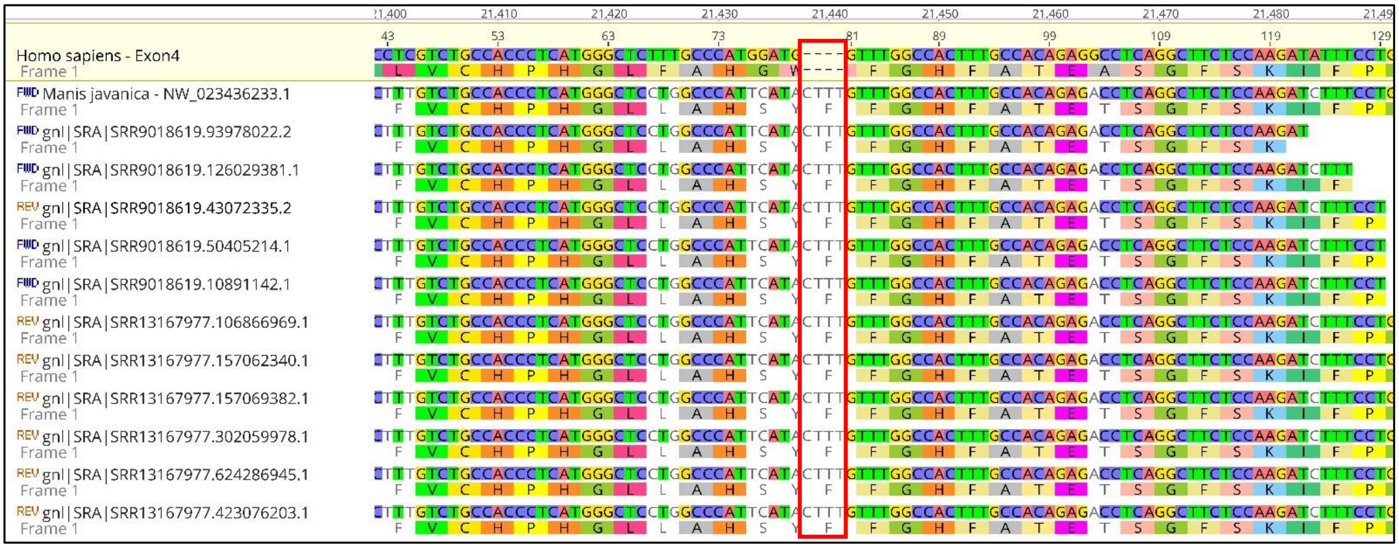

*Manis pentadactyla:*

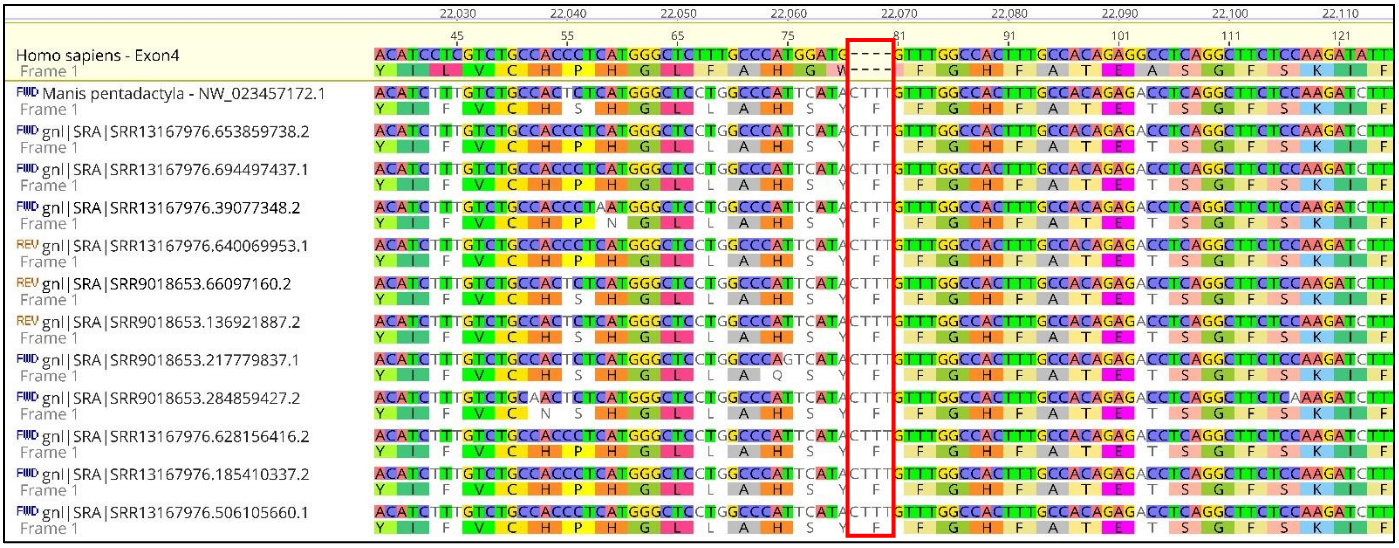

SRA Validation of a deletion in exon 4 of *Awat2* in *Phataginus tricuspis*:

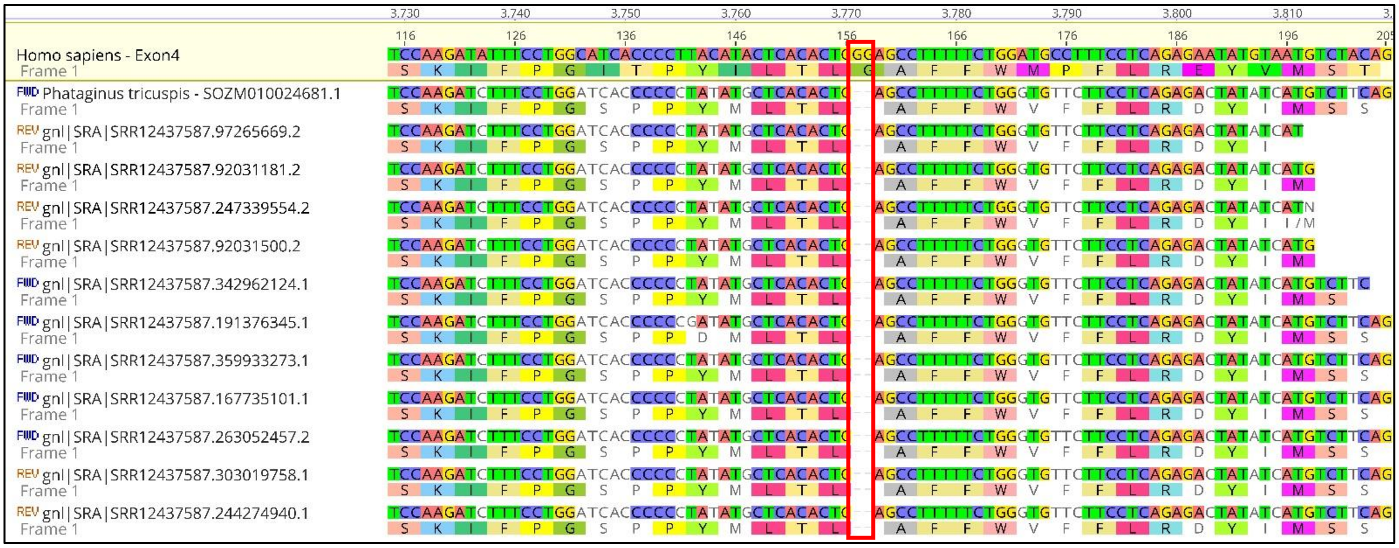

SRA Validation of a deletion in exon 2 of *Dgat2l6* in Manis:

*Manis javanica:*

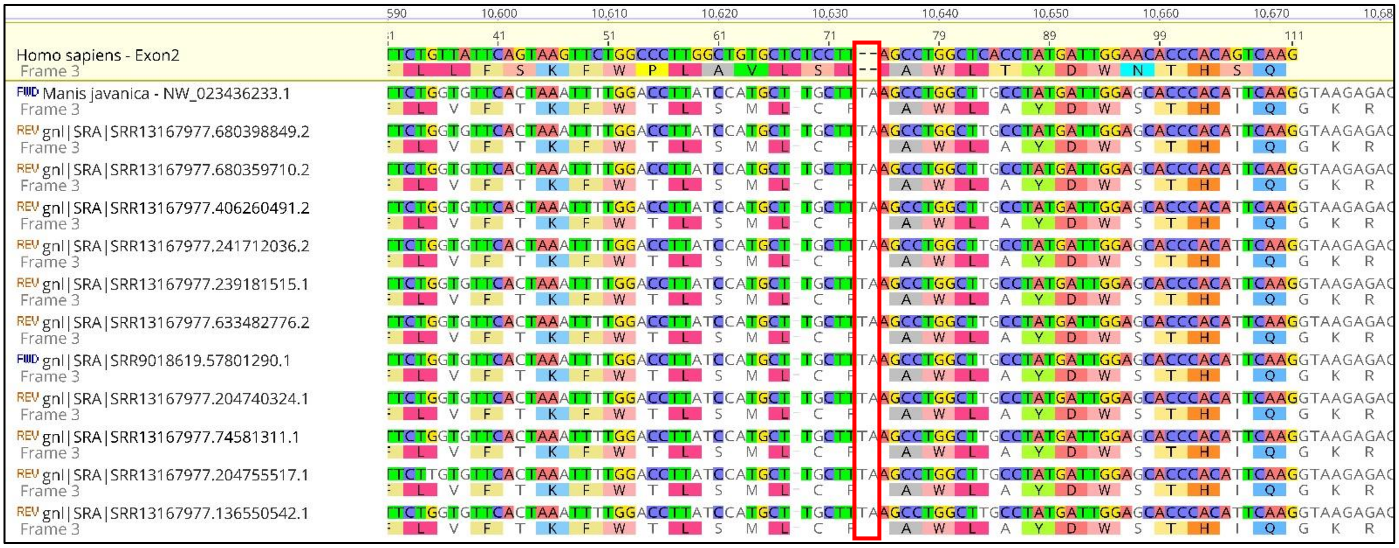

*Manis pentadactyla:*

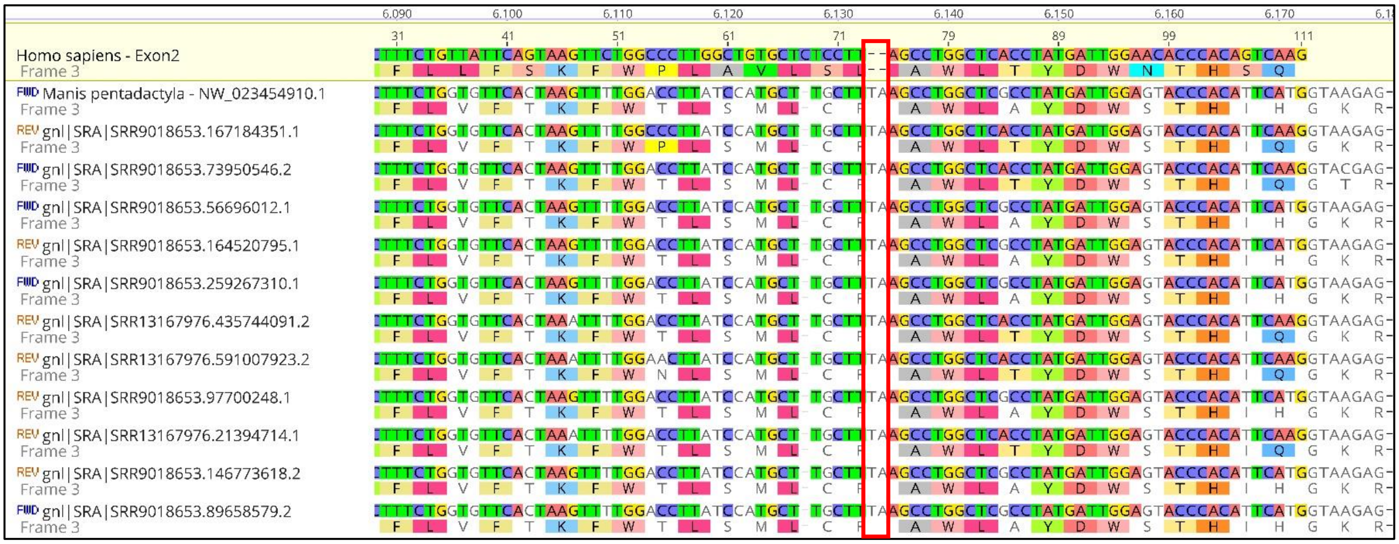

SRA Validation of an insertion in exon 2 of *Dgat2l6* in *Phataginus tricuspis*:

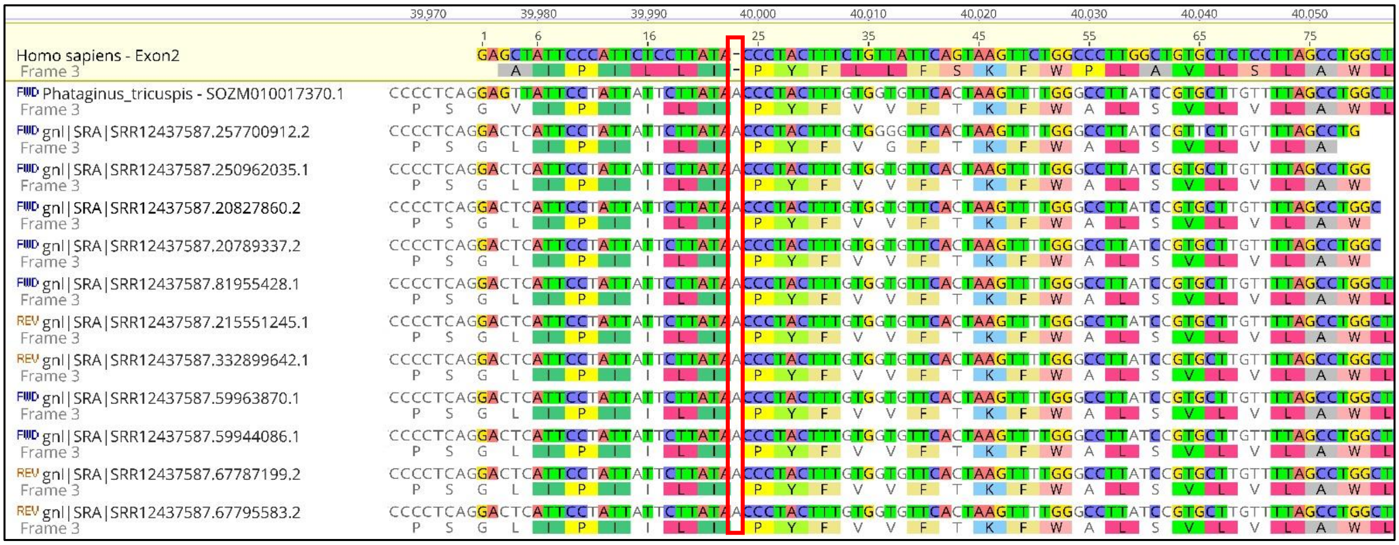

SRA Validation of a premature stop codon in exon 4 of *Eda2r* in Manis:

*Manis javanica:*

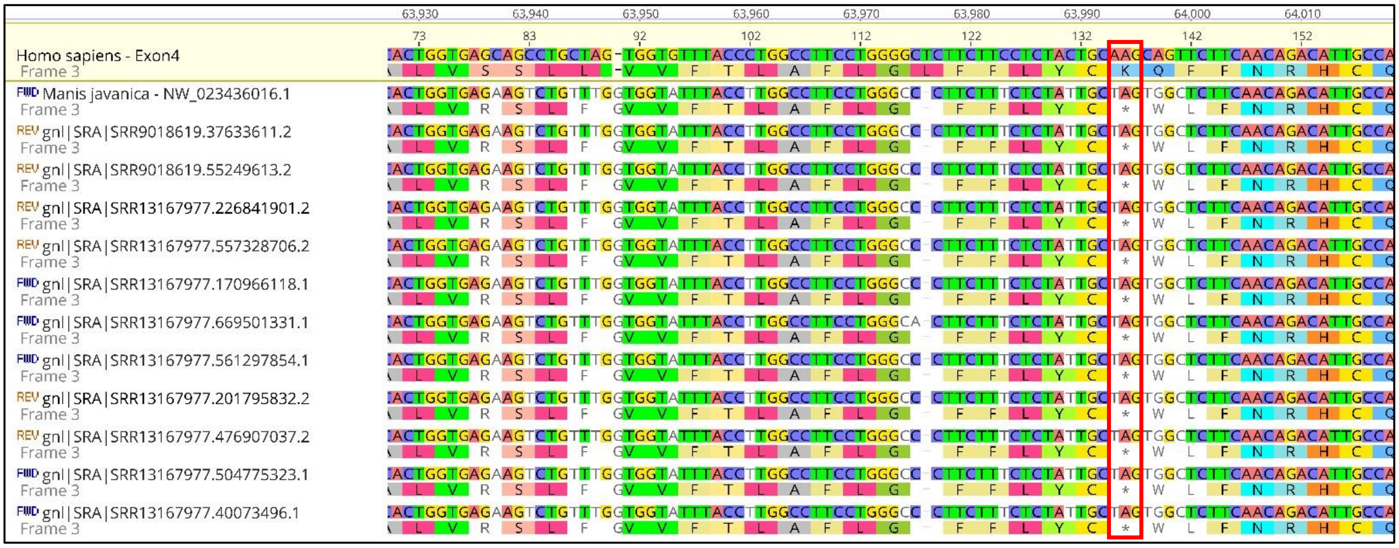


*Manis pentadactyla:*

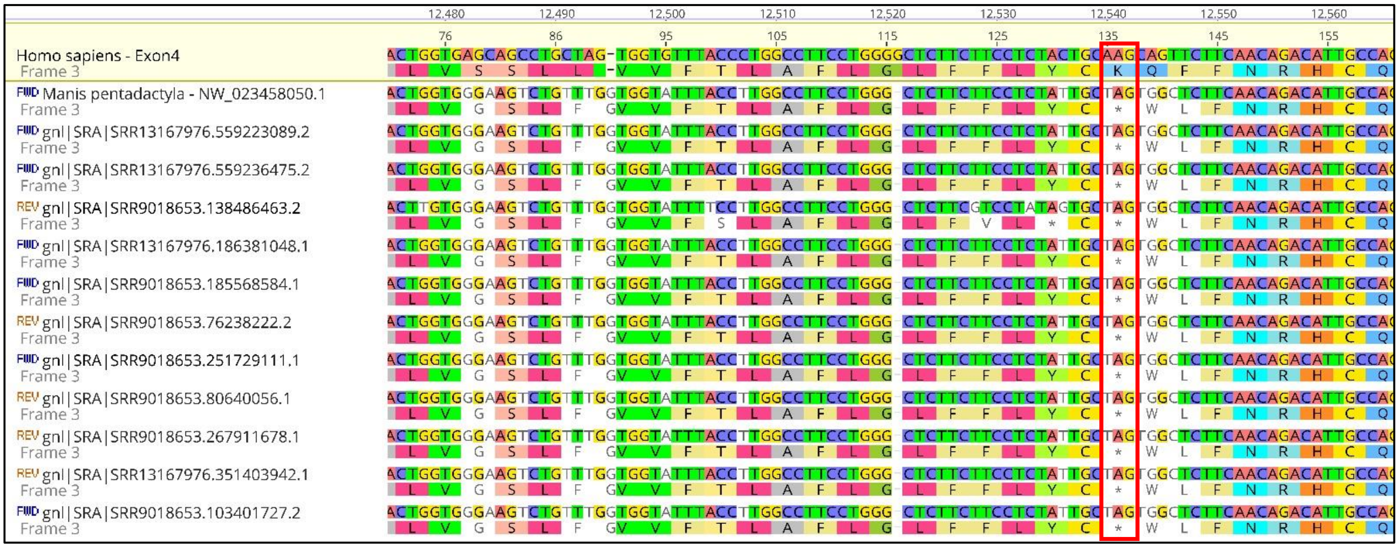

SRA Validation of an insertion in exon 3 of *Eda2r* in *Phataginus tricuspis:*

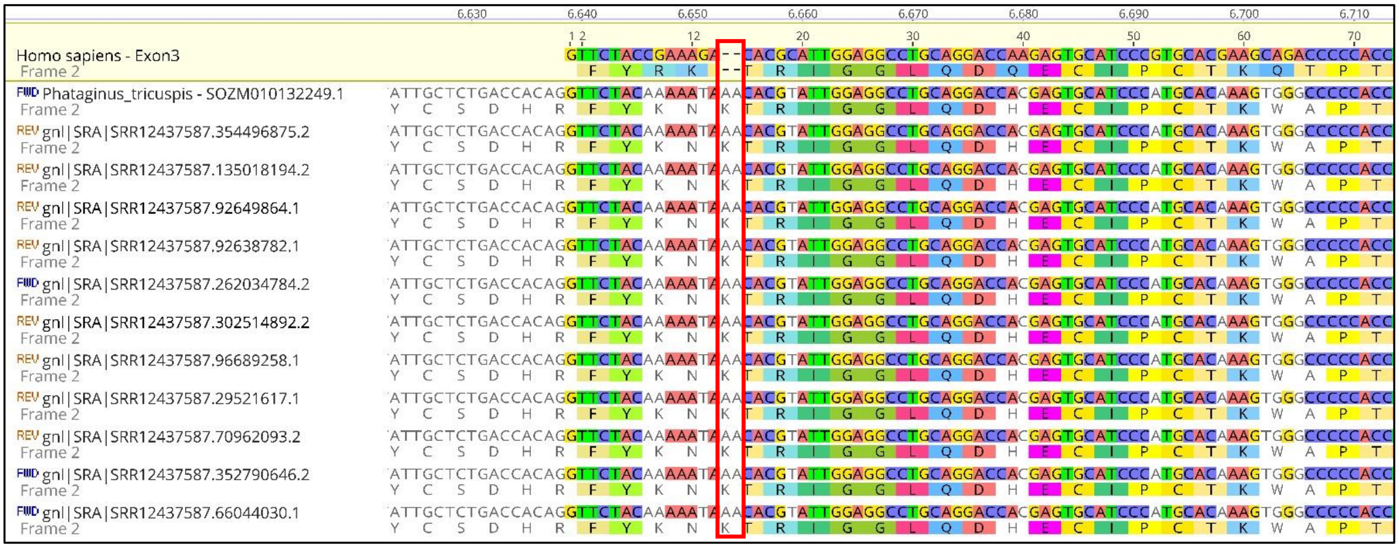


SRA Validation of a deletion in exon 2 of *Fabp9* in Manis:

*Manis javanica:*

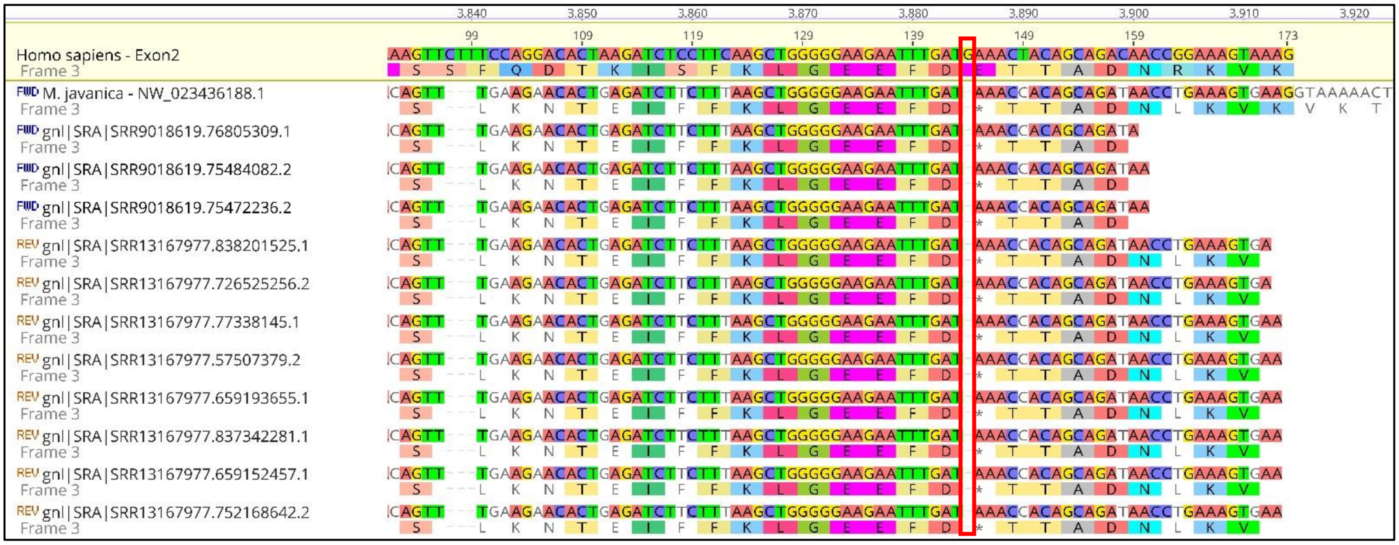

*Manis pentadactyla:*

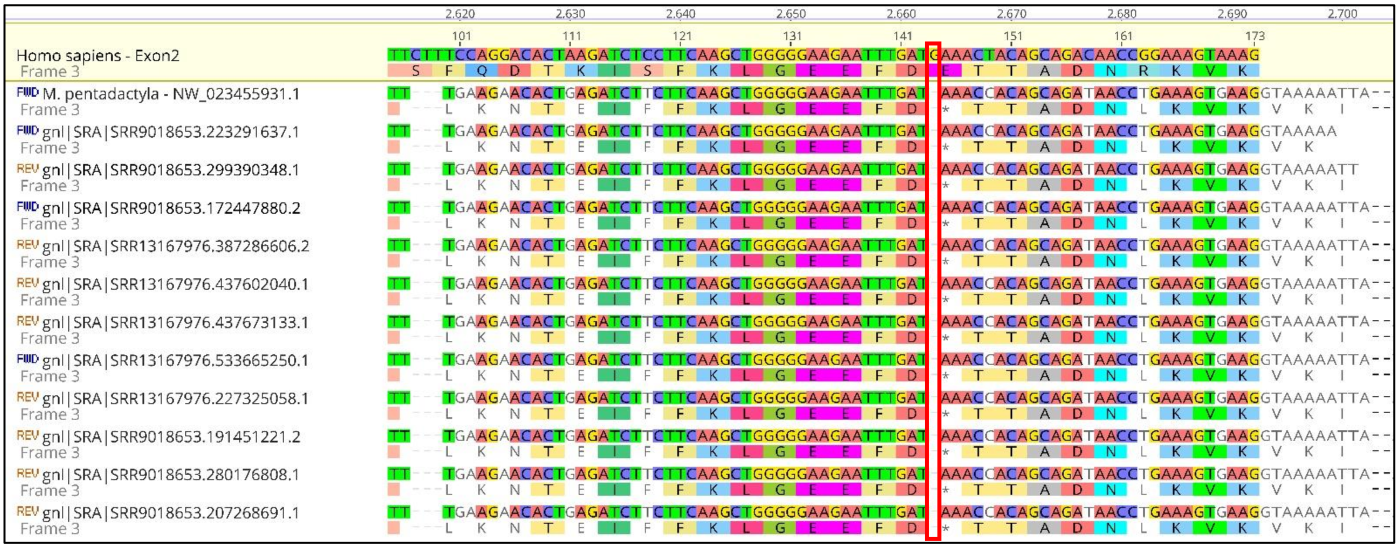


SRA Validation of a deletion in exon 1 of *Fabp9* in *Phataginus tricuspis:*

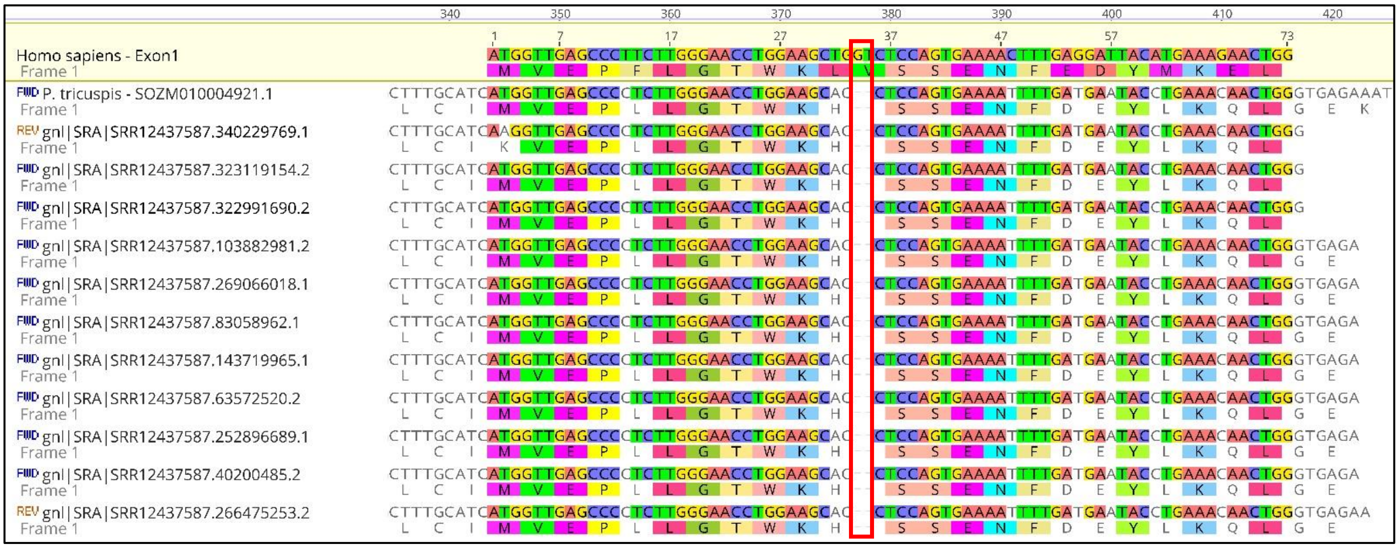

SRA Validation of a deletion in exon 3 of *Gsdmb* in Manis:

*Manis javanica:*

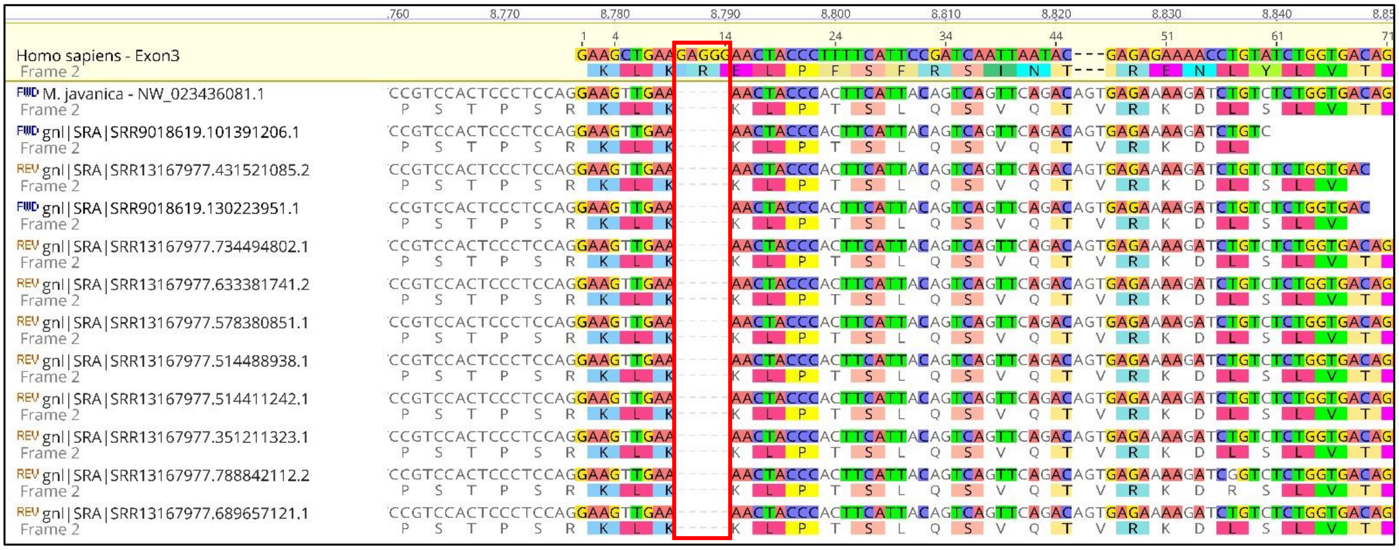

*Manis pentadactyla:*

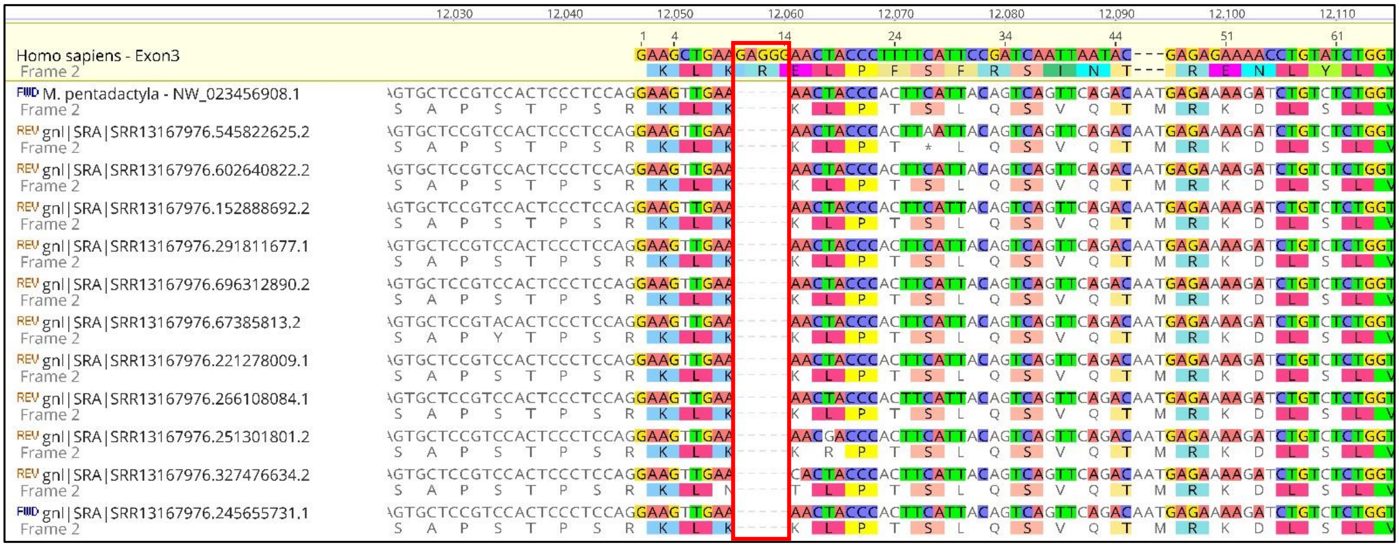

SRA Validation of a premature stop codon in exon 1 of *Gsdmb* in *Phataginus tricuspis:*

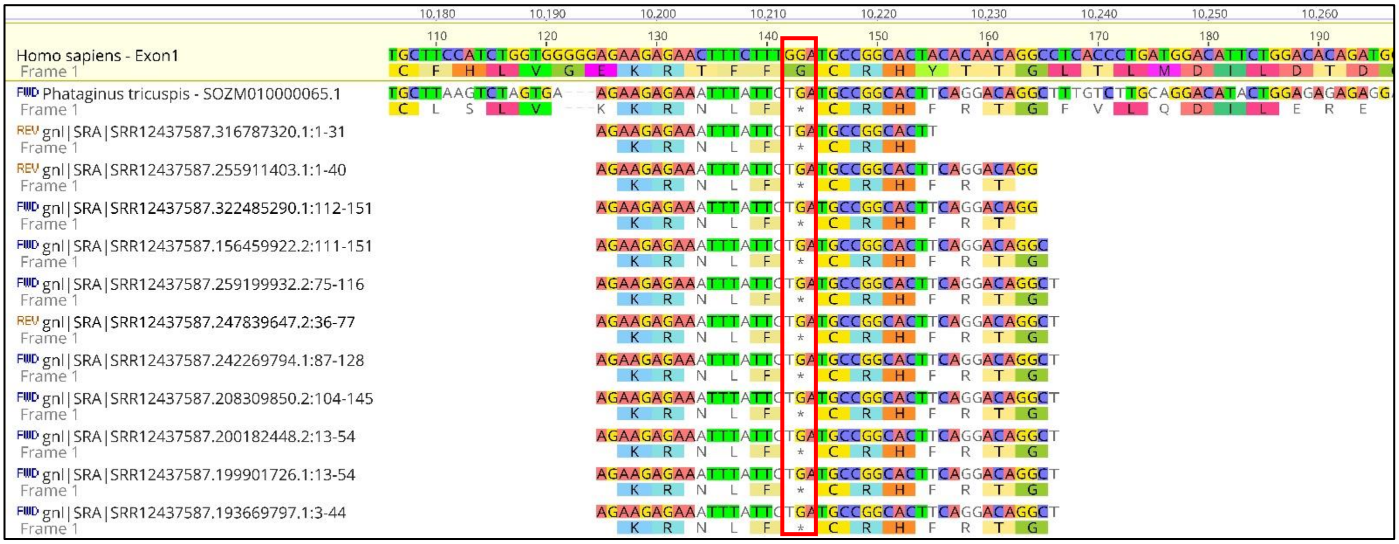


SRA Validation of a premature stop codon in exon 3 of *Mogat3* in *Manis javanica:*

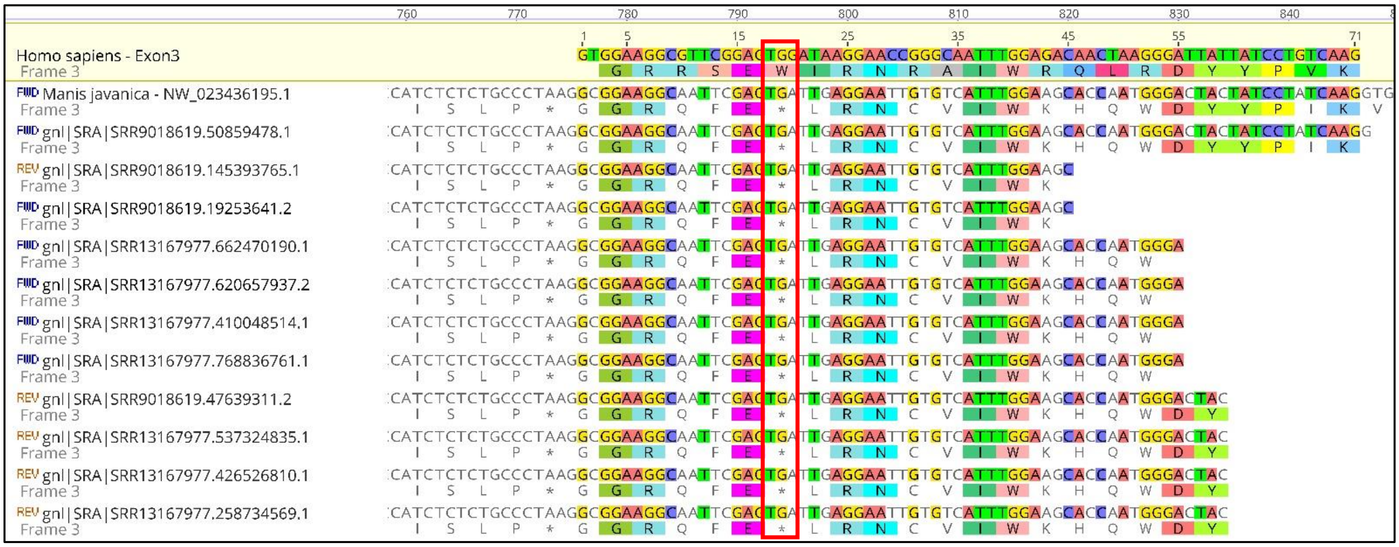

SRA Validation of a premature stop codon in exon 6 of *Mogat3* in *Manis pentadactyla:*

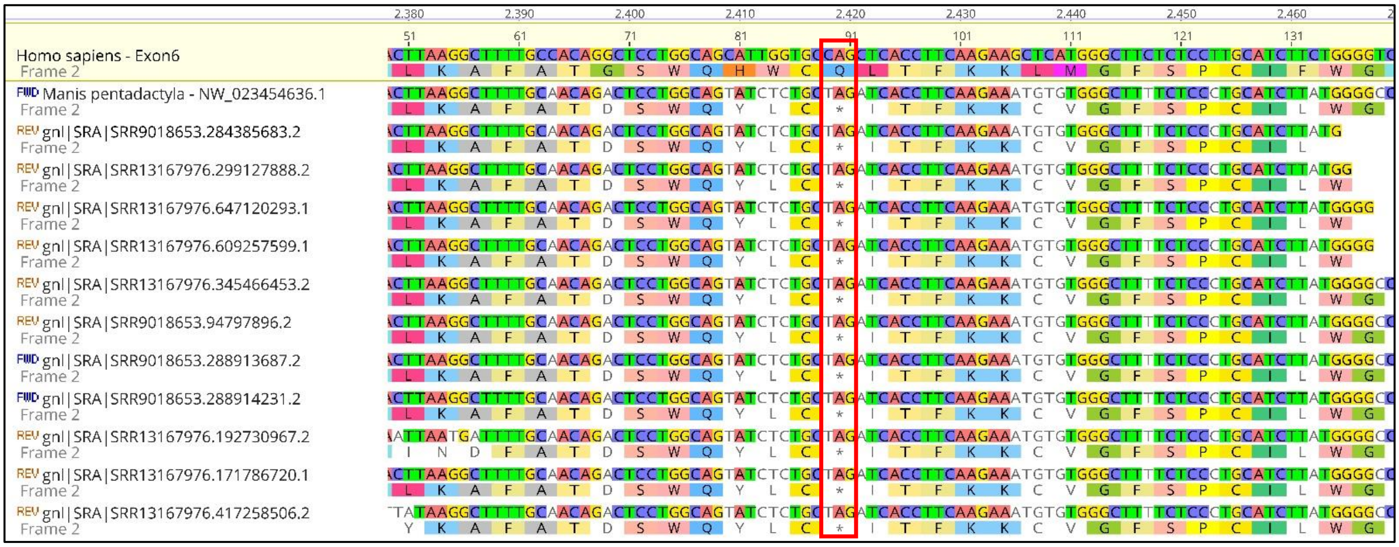

SRA Validation of a premature stop codon in exon 4 of *Mogat3* in *Phataginus tricuspis:*

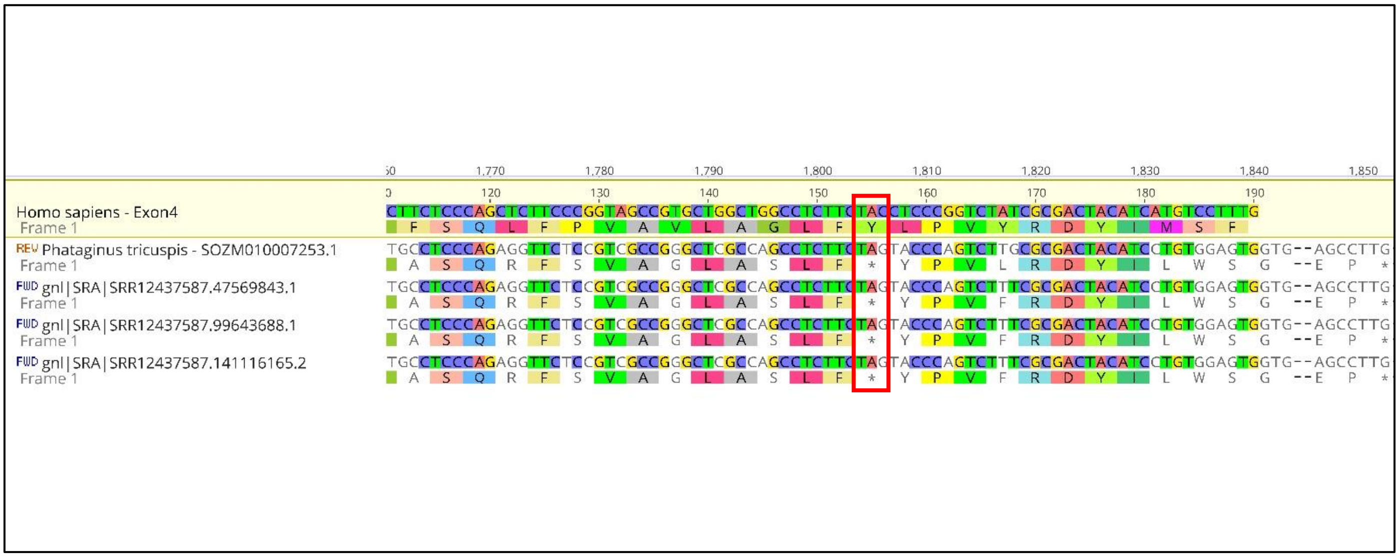

SRA Validation of a deletion in exon 1 of *Slurp1* in *Phataginus tricuspis:*

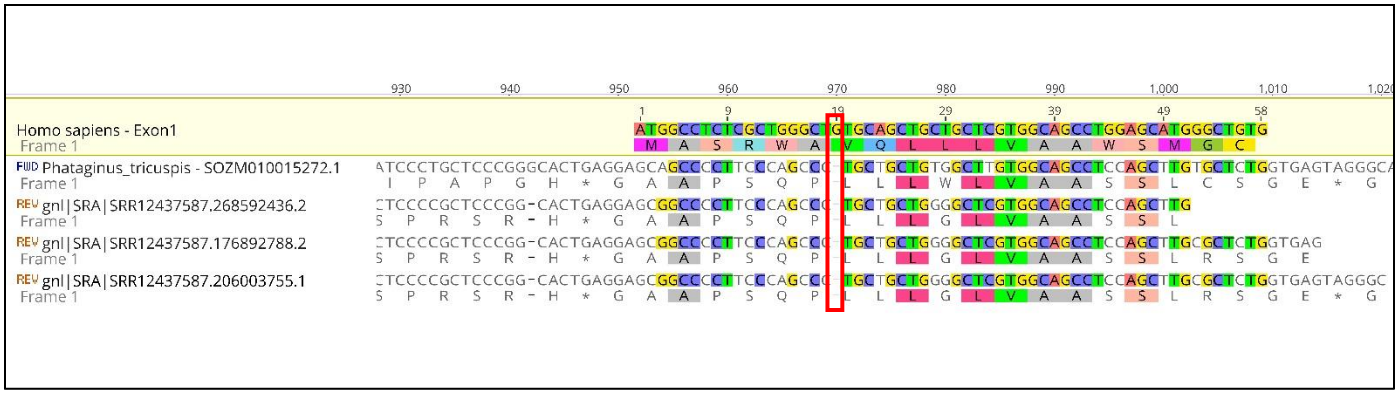


SRA Validation of a deletion in exon 2 of *Tchhl1* in Manis:

*Manis javanica:*

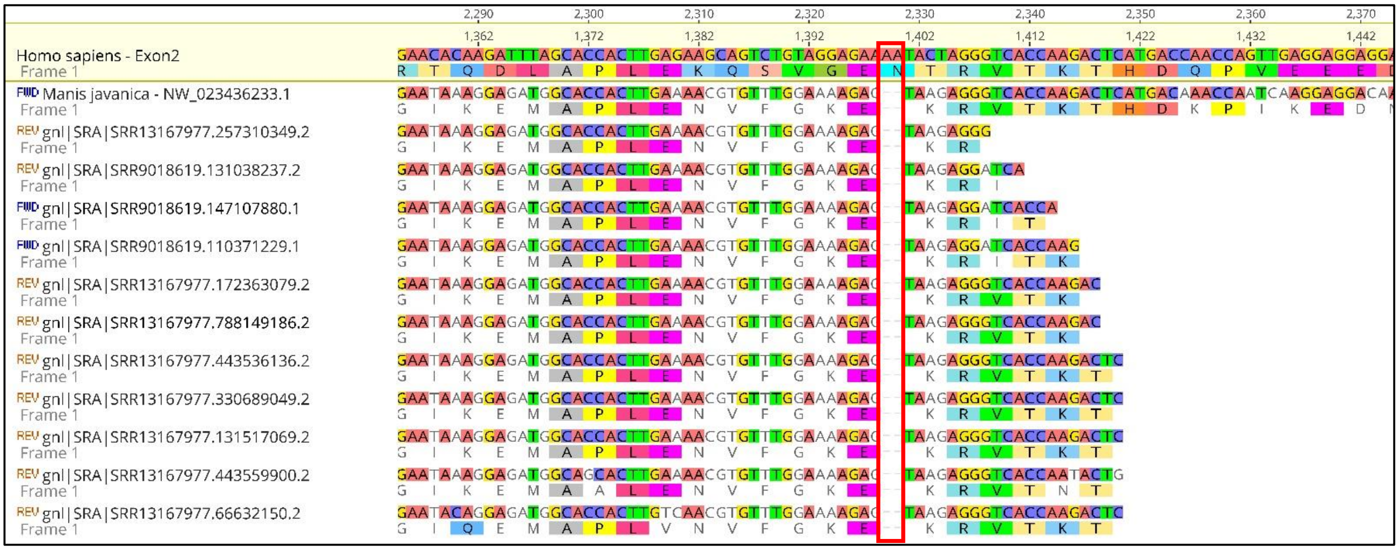

*Manis pentadactyla:*

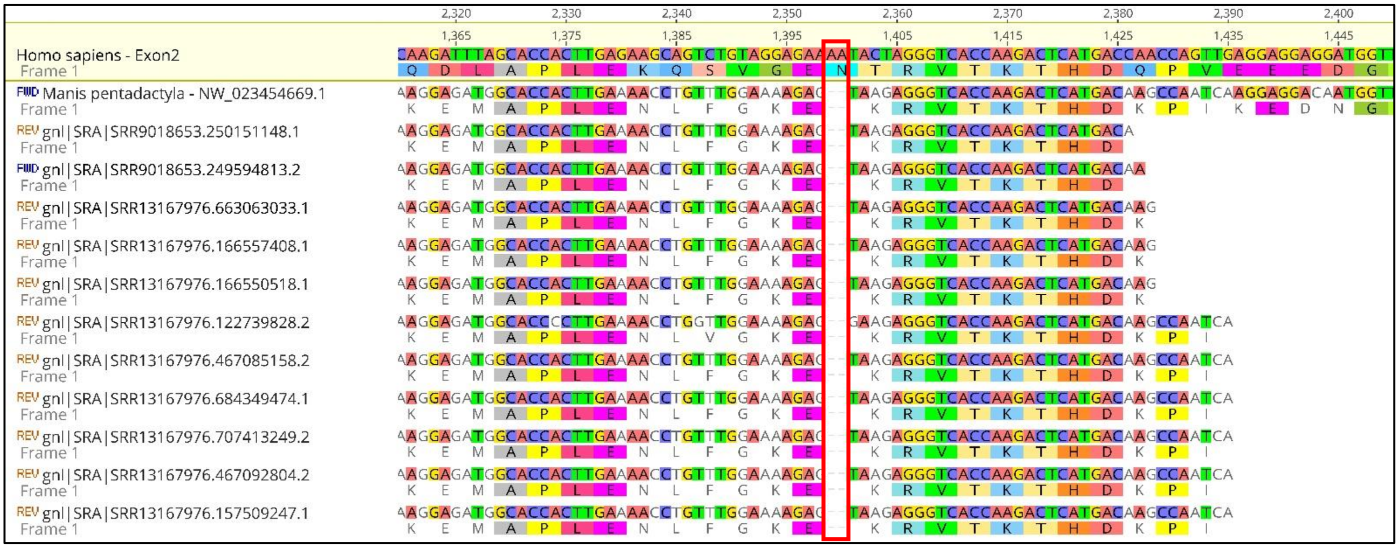
